# Stable Flow-induced Expression of KLK10 Inhibits Endothelial Inflammation and Atherosclerosis

**DOI:** 10.1101/2021.08.10.455857

**Authors:** Darian Williams, Marwa Mahmoud, Renfa Liu, Aitor Andueza, Sandeep Kumar, Dong-Won Kang, Jiahui Zhang, Ian Tamargo, Nicolas Villa-Roel, Kyung-In Baek, Hwakyoung Lee, Yongjin An, Leran Zhang, Edward W. Tate, Pritha Bagchi, Jan Pohl, Laurent O. Mosnier, Eleftherios P. Diamandis, Koichiro Mihara, Morley D. Hollenberg, Zhifei Dai, Hanjoong Jo

**Author notes:** Corresponding author: Hanjoong Jo, PhD, Wallace H. Coulter Distinguished Faculty Chair Professor, Wallace H. Coulter Department of Biomedical Engineering, Emory University and Georgia Institute of Technology, 1760 Haygood Drive, Health Science Research Building, E170, Atlanta, GA 30322, USA. DW, MM and RL equally contributed to this work.

## Abstract

**Introduction:** Atherosclerosis preferentially occurs in arterial regions exposed to disturbed blood flow (*d-flow*), while regions exposed to stable flow (*s-flow*) are protected. The proatherogenic and atheroprotective effects of *d-flow* and *s-flow* are mediated in part by the global changes in endothelial cell gene expression, which regulates endothelial dysfunction, inflammation, and atherosclerosis. Previously, we identified Kallikrein-Related Peptidase 10 (KLK10, a secreted serine protease) as a flow-sensitive gene in arterial endothelial cells, but its role in endothelial biology and atherosclerosis was unknown.

**Methods and Results:** Here, we show that KLK10 is upregulated under *s-flow* conditions and downregulated under *d-flow* conditions using *in vivo* mouse models and *in vitro* studies with cultured endothelial cells (ECs). Single-cell RNA sequencing (scRNAseq) and scATAC sequencing (scATACseq) study using the partial carotid ligation mouse model showed flow-regulated KLK10 expression at the epigenomic and transcription levels. Functionally, KLK10 protected against *d-flow*-induced inflammation and permeability dysfunction in human artery ECs (HAECs). Further, treatment of mice *in vivo* with rKLK10 decreased arterial endothelial inflammation in *d-flow* regions. Additionally, rKLK10 injection or ultrasound-mediated transfection of KLK10-expressing plasmids inhibited atherosclerosis in *ApoE*^-/-^ mice. Studies using pharmacological inhibitors and siRNAs revealed that the anti-inflammatory effects of KLK10 were mediated by a Protease Activated Receptors (PAR1/2)-dependent manner. However, unexpectedly, KLK10 did not cleave the PARs. Through a proteomics study, we identified HTRA1 (High-temperature requirement A serine peptidase 1), which bound and cleaved KLK10. Further, siRNA knockdown of HTRA1 prevented KLK10’s anti-inflammatory and barrier protective function in HAECs, suggesting that HTRA1 regulates KLK10 function. Moreover, KLK10 expression was significantly reduced in human coronary arteries with advanced atherosclerotic plaques compared to those with less severe plaques.

**Conclusion:** KLK10 is a flow-sensitive endothelial protein and, in collaboration with HTRA1, serves as an anti-inflammatory, barrier-protective, and anti-atherogenic factor.

## Introduction

Atherosclerosis is an inflammatory disease that preferentially occurs in branched or curved arterial regions exposed to disturbed flow (*d-flow),* while areas of stable flow (*s-flow*) are protected from atherosclerosis^1–4^. Endothelial cells (ECs) are equipped with several mechanosensors located at the luminal and abluminal surface, cell-cell junction, and cytoskeleton, which detect fluid shear stress and trigger cascades of signaling pathways and cellular responses^3–10^. *D-flow* induces endothelial dysfunction and atherosclerosis in large part by regulating flow-sensitive coding and non-coding genes, as wells as epigenetic modifiers^2, 11–13^. Using the partial carotid ligation (PCL) mouse model of atherosclerosis and transcriptomic studies, we identified hundreds of flow-sensitive genes in endothelial cells (ECs) that change by *d-flow* in the left carotid artery (LCA) compared to the *s-flow* in the right carotid artery (RCA)^14, 15^. Among the flow-sensitive genes, Kallikrein related-peptidase 10 (*KLK10)* was identified as one of the most flow-sensitive; with high expression under *s-flow* and low expression under *d-flow* conditions^15^. However, its role in endothelial function and atherosclerosis was not known.

KLK10 was initially identified as a normal epithelial cell-specific 1 (NES1) protein^16^ and is a member of the kallikrein-related peptidase ‘KLK’ family of 15 secreted serine proteases, which are found as a gene cluster on human chromosome (19q13.4)^17^. The KLKs are distinct from plasma kallikrein, which is encoded on a separate chromosome (4q35)^18^. Despite the chromosomal clustering of the KLKs, each enzyme has a unique tissue expression pattern with different cellular functions. Typically, the KLKs are produced as inactive full-length pre-propeptides, which are then secreted and activated by a complex process to yield active extracellular enzyme^18^. KLKs are involved in a wide variety of processes ranging from skin desquamation to tooth development, hypertension and cancer^19,20–24^.

KLK10 was initially discovered as a potential tumor suppressor with its expression downregulated in breast, prostate, testicular, and lung cancer^25–29^. Further studies, however, showed a more complex story as KLK10 is overexpressed in ovarian, pancreatic, and uterine cancer^30–33,34, 35^. However, the role of KLK10 for endothelial function and atherosclerosis is not known.

Here, we tested the hypothesis that KLK10 mediates the anti-atherogenic effects of *s-flow*, while the loss of KLK10 under *d-flow* conditions leads to pro-atherogenic effects. We also show a novel non-canonical mechanism by which HTRA1 (High-temperature requirement A serine peptidase 1) binds and regulates KLK10 function.

## Results

### KLK10 expression is increased by *s-flow* and decreased by *d-flow* in ECs *in vitro and in vivo*

We first validated our previous mouse gene array data at the mRNA and protein levels by additional qPCR, immunostaining, Western blots, and ELISA in ECs *in vivo* and *in vitro*. To validate the flow-dependent regulation of KLK10 expression *in vivo*, mouse PCL surgery was performed to induce *d-flow* in the left carotid artery (LCA) while maintaining *s-flow* in right carotid artery (RCA) (Figure 1a-d). Consistent with our previous data^14, 15^, KLK10 protein and mRNA expression was significantly higher in ECs in the *s-flow* RCA compared to the *d-flow* LCA (Figure 1b-d). Furthermore, KLK10 expression was observed only in the RCA endothelial layer but not in the medial layer, suggesting that KLK10 is expressed in ECs in the carotid artery region exposed to *s-flow*. In addition, KLK10 protein expression was reduced in the lesser curvature (LC; the athero-prone aortic arch region that is naturally and chronically exposed to *d-flow*) compared to the greater curvature region (GC; the athero-protected aortic arch region that is naturally and chronically exposed to *s-flow)* as shown by *en face* immunostaining (Figure 1e,f).

**Figure 1.**
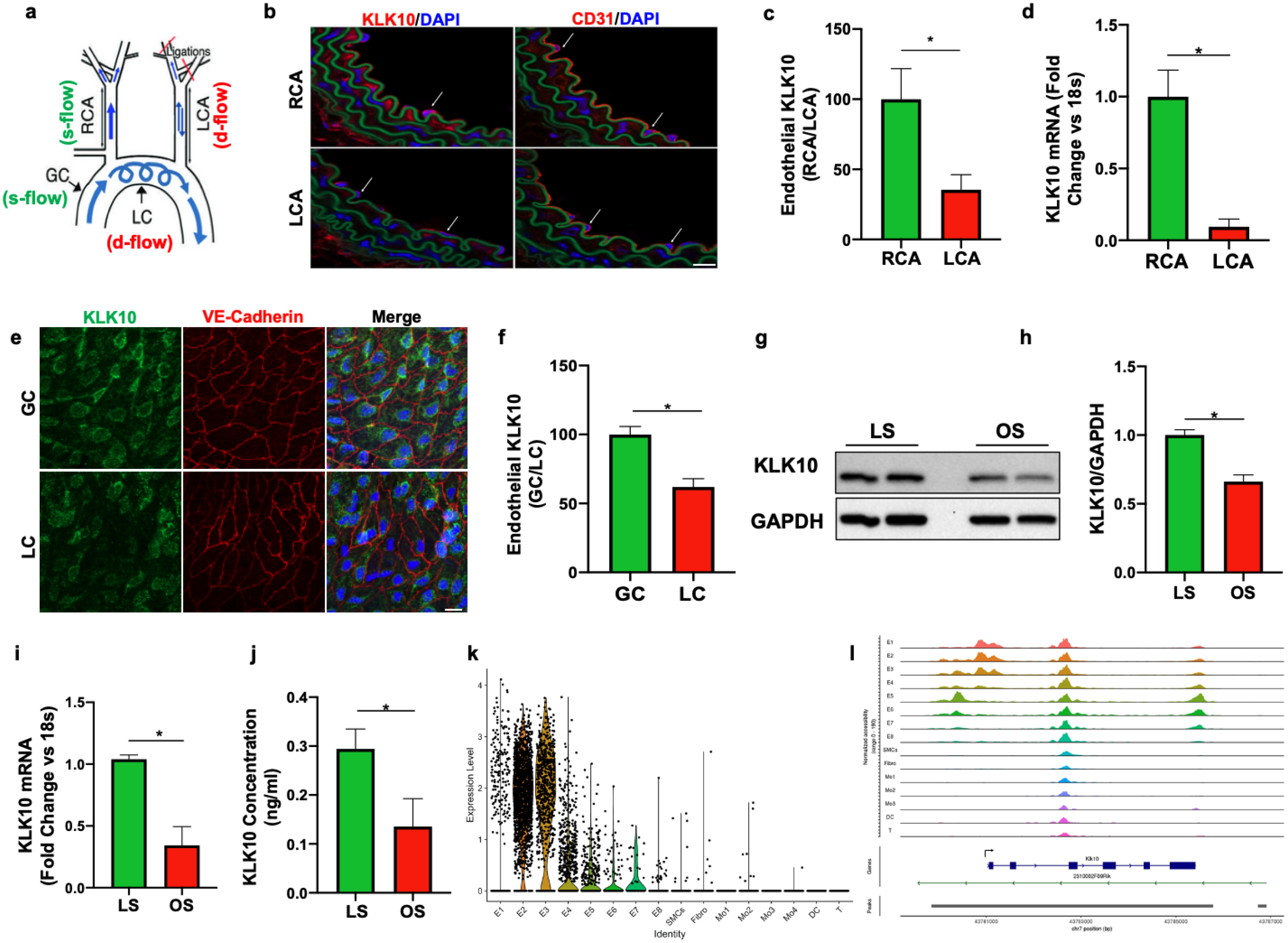
KLK10 expression is suppressed by disturbed flow (*d-flow*) and elevated by stable flow (s-flow) in endothelial cells *in vitro* and *in vivo.* (a) Depiction of the partial carotid ligation (PCL) surgery and flow-sensitive regions in the aortic arch: right carotid artery (RCA; *s-flow*), left carotid artery (LCA; *d-flow*), greater curvature (GC: *s-flow*), and lesser curvature (LC; *d-flow*). Two days following the PCL of C57BL/6 mice, RCA and LCA were collected to prepare for frozen section imaging (b-c) and (d) endothelial-enriched RNA preparation used for KLK10 qPCR. Shown are confocal images of immunostaining with KLK10 or CD31 antibodies (red) and counterstained with DAPI (blue) Scale bar=200 μm. Arrows indicate endothelial cells (c). Quantification of endothelial KLK10 fluorescence normalized to the % of the RCA. (e) Confocal images of e*n-face* immunostaining of the LC and GC with anti-KLK10 (green) and anti-VE-Cadherin (red) antibody are shown. Counterstained with DAPI (blue). Scale bar=20 μm. (f) Quantification of endothelial KLK10 fluorescence normalized to the % of the GC. (g-j) HAECs subjected to 24hr of unidirectional laminar shear (LS; 15 dynes/cm^2^) or oscillatory shear (OS; ±5 dynes/cm^2^) were used to measure expression of KLK10 mRNA by qPCR (g), KLK10 protein in cell lysates by western blot (h,i) and (j) KLK10 protein secreted to the conditioned media by ELISA. (k) Single-cell RNA sequencing (scRNAseq) and (l) single-cell ATAC sequencing (scATACseq) data sets (Andueza et al. Cell Reports) were re-analyzed. We found a flow-dependent response to KLK10 mRNA expression and chromatin accessibility, respectively, in ECs in the RCAs and LCAs at 2 days and 2 weeks after the PCL. KLK10 expression in ECs (E1-E8), smooth muscle cells (SMCs), fibroblasts (Fibro), monocytes/macrophages (Mo1-4), dendritic (DC), and T cells (T) are shown as we previously reported (Aitor Cell Reports). E2 and E3 represent ECs exposed to *s-flow* conditions, while E5 and E7 clusters represent ECs exposed to acute (2 days) *d-flow.* E6 and E8 represent ECs exposed to chronic (2 weeks) *d-flow.* (l) All data is represented as Mean±SEM, Paired two-tailed t-Test. n=4-6.*P≤0.05.

We next tested whether flow can regulate KLK10 expression *in vitro* using human aortic ECs (HAECs) exposed to unidirectional laminar shear (LS at 15 dynes/cm^2^) or oscillatory shear (OS at ±5 dynes/cm^2^ at 1 Hz) for 24h using the cone and plate viscometer, mimicking *s-flow* and *d-flow* conditions *in vivo*, respectively^36, 37^. KLK10 mRNA (Figure 1g), protein in cell lysates (Figure 1h,i), and secreted protein in the conditioned media were decreased by OS and increased by LS (Figure 1j), confirming the role of KLK10 as a flow-sensitive gene and protein *in vivo* and *in vitro*.

We further confirmed the flow-dependent expression of KLK10 by re-analyzing the single-cell RNA sequencing (scRNAseq) and ATAC sequencing (scATACseq) data sets that we recently published using the PCL model^38^. For the scRNAseq and scATACseq study, single-cells and -nuclei obtained from the LCAs and RCAs, respectively, at 2 days or 2 weeks after the PCL were used. As described previously, the carotid artery wall cells were identified as EC clusters (E1-E8), smooth muscle cells (SMC), fibroblasts (Fibro), monocytes/macrophages (Mo1-4), dendritic cells (DC) and T-cells^38^ (Figure 1K and I). E2 and E3 represent athero-protected endothelial populations exposed to *s-flow* in the RCA, whereas E5 and E7 represent ECs exposed to acute *d-flow* (for 2 days). E6 and E8 populations represent athero-prone ECs exposed to chronic *d-flow* in the LCA (2 weeks). As shown in the scRNAseq data analysis (Figure 1K), KLK10 expression is highest in *s-flow* (E2 and E3) and decreases in response to acute (E5 and E7) and chronic (E6 and E8) *d-flow*. It also shows KLK10 expression is specific to ECs and not expressed in other cell types studied in the carotid. Similarly, scATACseq data (Figure 1I) showed that the KLK10 promoter region is open and accessible (indicating active transcription status) in *s-flow* conditions but closed and inaccessible (indicating inactive transcription status) under *d-flow* conditions. Together, both the scRNAseq and scATACseq results demonstrate that KLK10 expression is potently regulated by flow at the epigenomic and transcriptome level, supporting the *in vitro* and *in vivo* results shown above (Figure 1b-g).

### KLK10 regulates the endothelial inflammation and permeability

We next tested if KLK10 regulates EC function by evaluating its role in endothelial inflammatory response, tube formation, migration, proliferation, and apoptosis, which play critical roles in the pathogenesis of atherosclerosis. Treatment of human umbilical vein ECs (HUVECs) with rKLK10 significantly inhibited migration and tube formation, but not proliferation and apoptosis (Supplementary Figure S1). In addition, transfection of HAECs with plasmids to overexpress KLK10 reduced THP1 monocyte adhesion to the ECs in response to TNFα and under basal conditions (Figure 2a; Figure S2). Next, we pre-treated HAECs overnight with increasing concentrations of rKLK10, followed by TNFα treatment (5 ng/mL for 4h). Treatment with rKLK10 significantly inhibited monocyte adhesion to ECs in a concentration-dependent manner (Figure 2b). Of note, the anti-inflammatory effect of rKLK10 was lost if rKLK10 was heated, implicating the importance of the enzymatic activity or native conformation of KLK10 (Figure 2b, Figure S4). Furthermore, treatment with rKLK10 significantly inhibited mRNA and protein expression of the pro-inflammatory adhesion molecules VCAM1 and ICAM1 (Figure 2c-g), as well as the phosphorylation and nuclear localization of p65 in response to TNFα (Figure S5).

**Figure. 2.**
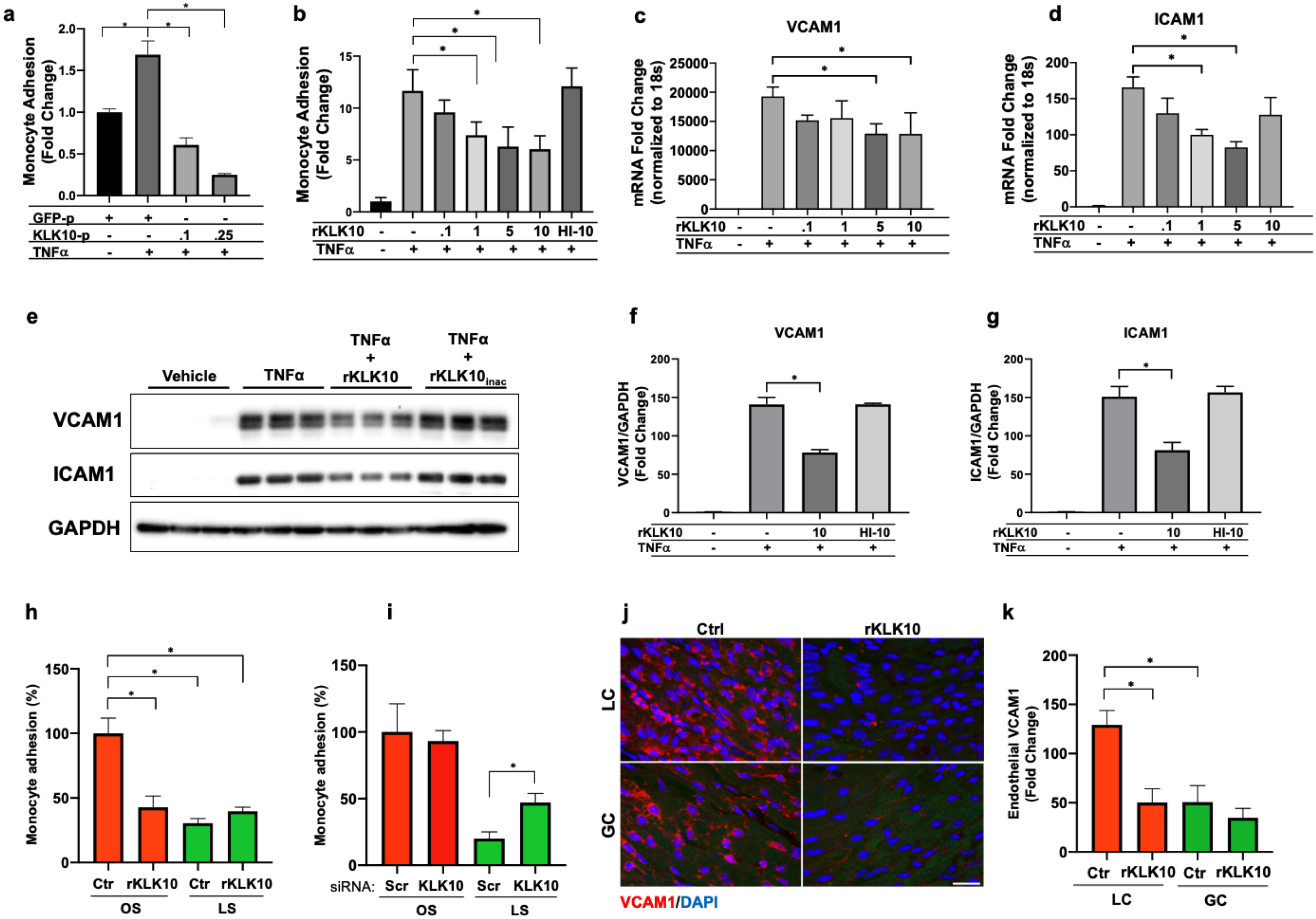
KLK10 inhibits inflammation in endothelial cells *in vitro* and *in vivo*. (a) THP1 monocyte adhesion assay was carried out in HAECs transfected with 0.1μg or 0.25μg of KLK10 plasmid (KLK10-p) or GFP plasmid (GFP-p) for 48hrs followed by TNFɑ treatment (5 ng/mL for 4hr). Data is represented as fold change in monocyte adhesion normalized to GFP-p control. (b) THP1 monocyte adhesion assay was carried out in HAECs treated with rKLK10 (0.1-10 ng/mL) or heat-inactivated rKLK10 (HI-10) for 24hrs followed by TNFɑ treatment (5 ng/mL for 4hr). Data is represented as fold change in monocyte adhesion normalized to vehicle control. (c-g) HAECs were treated with rKLK10 (0.1-10 ng/mL for 24hrs) followed by TNFɑ treatment (5 ng/mL for 4hr) and expression of inflammatory marker VCAM1 and ICAM1 expression were assessed by qPCR (c,d) or western blot (e-g). Data is normalized to housekeeper genes 18s and GAPDH and represented as fold change of the vehicle control. (h,i) Monocyte adhesion assay was conducted on HAECs subjected to 24hr of either LS (15 dynes/cm^2^) or OS (±5 dynes/cm^2^) with either (h) rKLK10 (100 ng/mL) or (i) KLK10 siRNA (50 nM) for KLK10 or a non-targeting siRNA control. Data is represented as % of monocyte adhesion normalized to the control OS condition. (j) C57BL/6 mice were injected with rKLK10 (0.6 mg/kg) or a vehicle control on days 0, 2, and 4, and sacrificed on day 5. The aortic arches were *en-face* immunostained using an anti-VCAM1 antibody (red) and Dapi (blue). (k) Quantification of endothelial VCAM1 expression represented as fold change normalized to control LC condition. Scale bar=10 μm. All data is represented as Mean±SEM, One-way ANOVA with Bonferroni correction for multiple comparisons. Mean±SEM n=4-6.*P≤0.05.

We then tested the effect of KLK10 on the endothelial inflammatory response as measured by monocyte adhesion under flow conditions *in vitro* and *in vivo*. rKLK10 treatment inhibited OS-induced monocyte adhesion in HAECs (Figure 2h). In contrast, siRNA-mediated knockdown of KLK10 significantly increased monocyte adhesion under LS conditions (Figure 2i; Figure S3). We next tested if rKLK10 could affect the endothelial inflammation in naturally flow-disturbed LC of the aortic arch in mice. Treatment with rKLK10 *in vivo* (intravenous injection every 2 days for 5 days at 0.6 mg/kg) dramatically reduced VCAM1 expression in the *d-flow* (LC) region in the aortic arch of these mice (Figure 2j,k). We also observed a dose-dependent effect of rKLK10 on VCAM1 expression (Figure S6). These results demonstrate that either KLK10 overexpression using plasmids or rKLK10 treatment protects against EC inflammation both *in vitro* and *in vivo* under TNFα or *d-flow* conditions, whereas the reduction of KLK10 under *d-flow* condition or KLK10 knockdown using siRNA increases inflammation.

Next, we tested if rKLK10 treatment can similarly protect the permeability barrier function of ECs. As a positive control, thrombin treatment increased the permeability of HAECs as measured by increased binding of fluorescently labeled (FITC)-Avidin to biotin-gelatin as reported previously^39^. Overnight rKLK10 pre-treatment prevented the permeability increase induced by thrombin in HAECs (Figure 3a,b). Similarly, rKLK10 reduced the permeability induced by OS (Figure 3c,d), further demonstrating the protective role of KLK10 in endothelial inflammation and barrier function.

**Figure 3.**
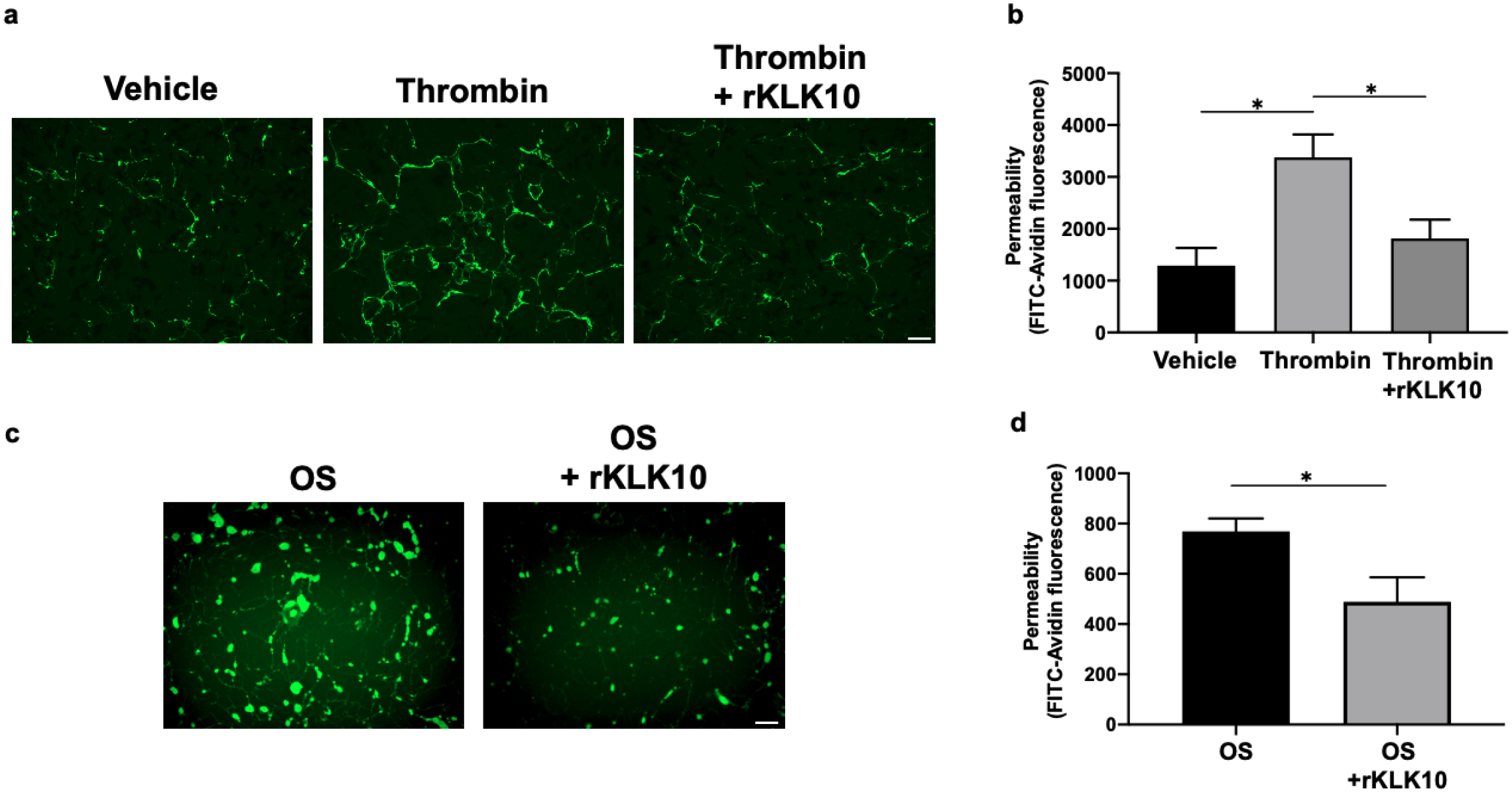
KLK10 protects endothelial permeability against thrombin and OS. (a) HAECs were grown to confluency on biotinylated gelatin and were treated with rKLK10 (10 ng/mL) or vehicle overnight followed by thrombin (5 U/mL for 30 minutes). Endothelial permeability was then measured by the binding of FITC-avidin to the biotinylated gelatin. (b) Quantification of endothelial permeability measured as FITC-Avidin fluorescence. (c) HAECs were grown to confluency on biotinylated gelatin and were subjected OS (±5 dynes/cm^2^) with rKLK10 (10 ng/mL) or vehicle for 20hrs. Endothelial permeability was then measured by the binding of FITC-avidin to the biotinylated gelatin. Scale bar=100 μm. All data is represented as Mean±SEM, n=4-6. One-way ANOVA with Bonferroni correction (b) or paired two-tailed T-Test (d).*P≤0.05.

### Treatment with rKLK10 inhibits atherosclerosis in *ApoE*^-/-^ mice

Given its anti-inflammatory and barrier protective effect in ECs, we tested if atherosclerosis development could be prevented by treating mice with rKLK10. For this study, we used the PCL model of atherosclerosis to induce atherosclerosis rapidly in a flow-dependent manner in hyperlipidemic *ApoE*^-/-^ mice fed with a high-fat diet. Injection with rKLK10 (twice per week at 0.6 mg/kg for 3 weeks post-PCL surgery) significantly reduced atherosclerosis development and macrophage accumulation in the LCA (Figure 4a-e). The rKLK10 treatment showed no effect on plasma levels of total, LDL(low-density lipoprotein), and HDL (high-density lipoprotein) cholesterols and triglycerides (Figure 4 f-i). Thus, rKLK10 showed an anti-atherogenic effect *in vivo*.

**Figure 4.**
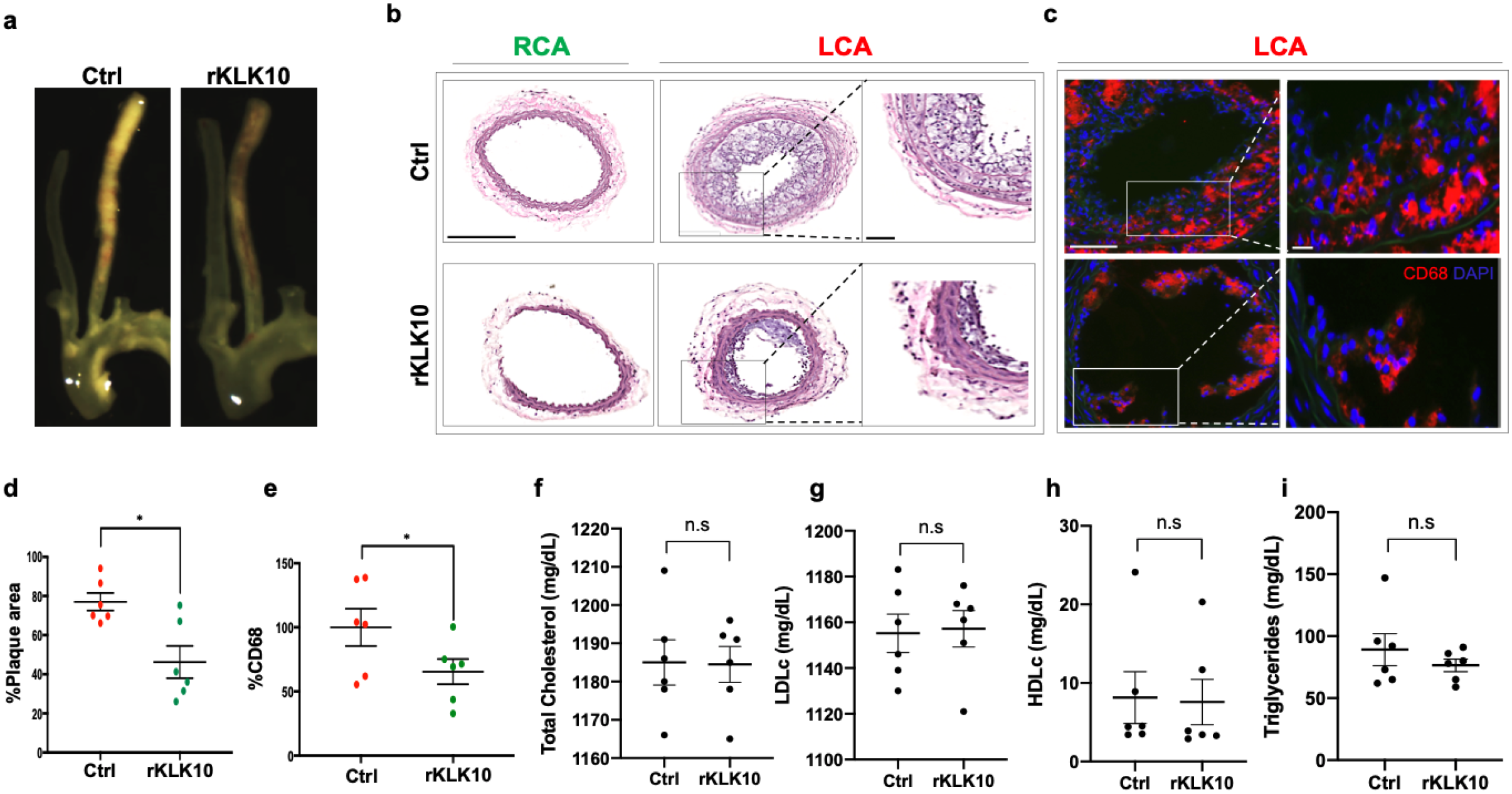
Treatment with rKLK10 inhibits atherosclerosis development in *ApoE^-/-^* mice. (a) *ApoE^-/-^* were subjected to partial carotid ligation and high fat diet feeding. The mice received either rKLK10 (0.6 mg/kg) or vehicle injection every three days for the duration of three weeks. LCA show plaque development, which was reduced by rKLK10 as shown by dissection microscopy. Frozen sections from the LCA and RCA were stained with (b) H&E and (c) for CD68 in LCA. DAPI (blue). Scale bar low mag=250 μm, high mag=50 μm. (d) Plaque area was quantified from H&E staining and is represented as the % plaque area. (e) CD68 macrophage accumulation was quantified and is represented as the CD68 % normalized to the control. (f-i) Plasma lipid analysis of (f) total cholesterol, (g) LDL cholesterol, (h) HDL cholesterol, or (i) triglycerides showed no effect of rKLK10 compared to control. All data is represented as Mean±SEM, *P≤0.05, Paired two-tailed t-test. N=6.

### Ultrasound-mediated overexpression of KLK10 inhibits atherosclerosis in *ApoE^-/-^* mice

We next asked whether overexpression of KLK10 using a plasmid vector could also inhibit atherosclerosis *in vivo*. For this study, we injected either KLK10 plasmid (pCMV-Igκ-KLK10-T2A-Luc) or luciferase plasmid (pCMV-Luc) as a control along with microbubbles to the hind-limbs of *ApoE*^-/-^ mice and sonoporated the legs with ultrasound as previously described^40–42^. The plasmid injection and sonoporation was repeated 10 days later to ensure sustained protein expression for the duration of the study. Bioluminescence imaging showed that all mice expressed luciferase in the hind-limbs at the conclusion of the study, indicating successful overexpression of the plasmids (Fig 5a).

**Figure 5.**
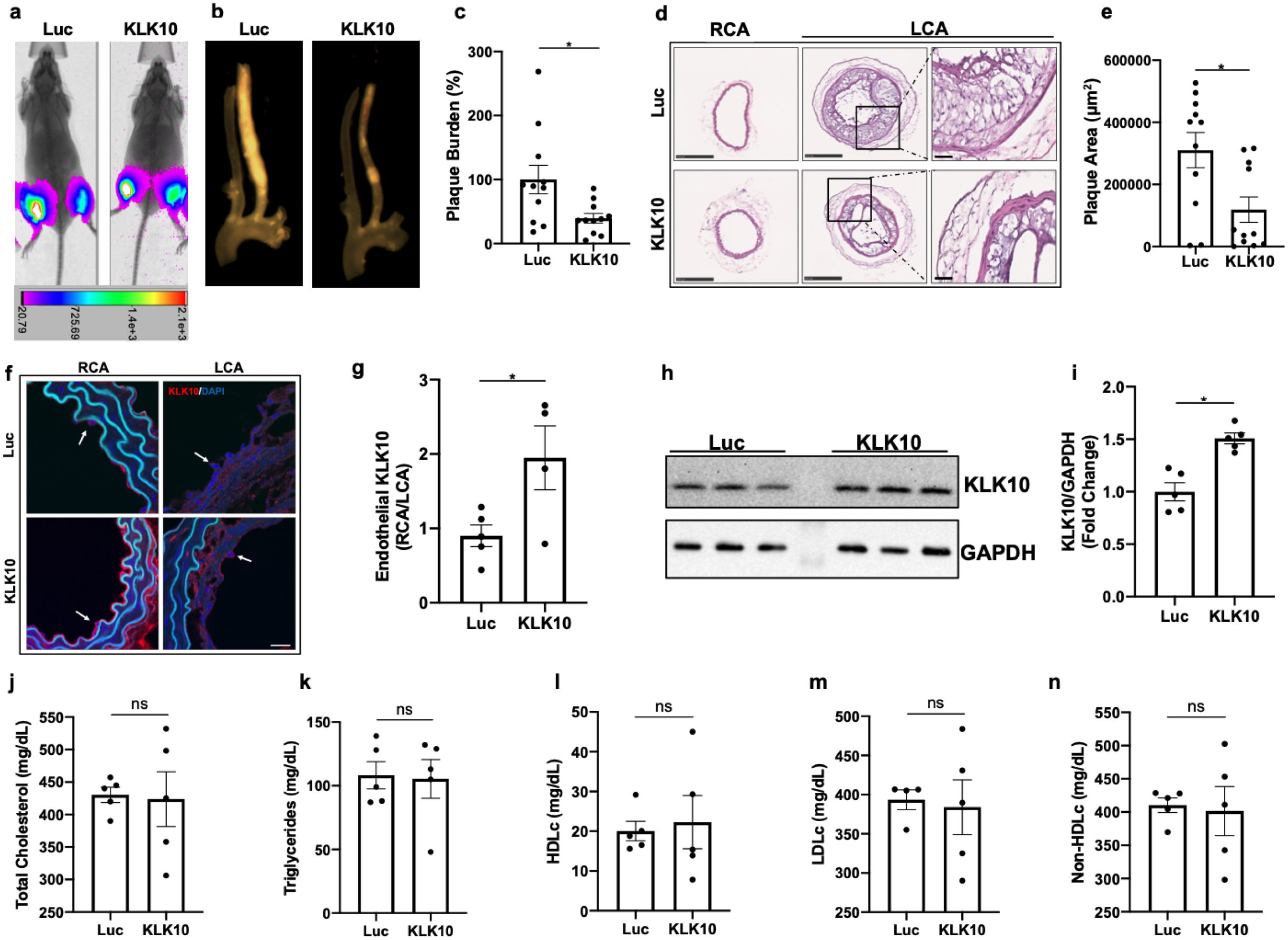
Ultrasound-mediated overexpression of KLK10 plasmid inhibits atherosclerosis development. (a) Bioluminescent imaging of *ApoE^-/-^* PCL mice on a high-fat diet injected with Luciferase control plasmid or KLK10-Luciferase plasmid, measured in photons/second. (b) Gross plaque images of excised carotid arteries and (c) quantification of plaque burden normalized to the % of the Luciferase control. (d) H&E staining of sections from the LCA and RCA of mice injected with Luciferase control plasmid or KLK10-Luciferase plasmid. Scale bar low mag=250 μm, high mag=50 μm. (e) Quantification of plaque area measured in μm^2^. (f) Immunostaining of sections from the RCA and LCA were further stained with anti-KLK10 (red) and DAPI (blue). Arrows indicate the endothelial layer. Scale bar=20 μm. (g) Quantification of endothelial KLK10 fluorescence represented as fold change normalized to Luciferase control. (h) Western blot assessing KLK10 expression in lung tissue from mice injected with control luciferase plasmid or KLK10 plasmid. GAPDH was used as a loading control. (i) Quantification of KLK10 expression in normalized to GAPDH and Luciferase control. (j-n) Plasma lipid analysis of (j) total cholesterol, k) triglycerides, (l) HDL cholesterol, (m) LDL cholesterol, or (n) non-HDL cholesterol. All data is represented as Mean±SEM, *P≤0.05, Paired two-tailed t-test. N=4-11.

Atherosclerotic plaque formation in the LCA was significantly reduced in the KLK10 overexpressing mice compared to the luciferase control (Fig 5b). Further assessment of the carotid artery sections by histochemical staining with hematoxylin and eosin (Figure 5d,e) showed decreased plaque area in the LCA of these mice. Circulating KLK10 levels in the mice measured by ELISA at the time of sacrifice showed no measurable difference between the luciferase and KLK10 groups (Figure S7). This may be due to a waning plasmid expression at the sacrifice time. However, we found higher levels of KLK10 staining at the endothelial layer in the LCA and RCA (Figure 5f,g), as well as in the lung tissue samples as shown by western blot (Figure 5h,i). These results demonstrate that treatment with KLK10 by either rKLK10 or KLK10 expression vector can inhibit atherosclerosis development in *ApoE*^-/-^ mice.

### KLK10 inhibits endothelial inflammation in a protease activated receptor-1/2 (PAR1/2)-dependent manner, but without cleaving their N-termini

PARs (1, 2 and 4) have been shown to mediate the effects of some KLKs (e.g. KLK5, 6, and 14)^43, 44^. Therefore, we tested if the anti-inflammatory effect of KLK10 could be mediated by PARs. Since our gene array and qPCR data showed that mouse artery ECs express PAR1 and PAR2, but not PAR3 and PAR4^14, 15^, we tested whether PAR1 and PAR2 mediate the KLK10 effect in ECs under OS condition by using two independent approaches - specific pharmacological inhibitors and siRNAs. ECs were first exposed to OS for 16h to induce an endothelial inflammatory response and treated with either the PAR1 inhibitor (SCH79797, 0.1μM) or PAR2 inhibitor (FSLLRY-NH2, 10μM). Both PAR inhibitors prevented the anti-inflammatory effect of KLK10 (Figure 6a), suggesting the role of PAR1/2 in mediating the KLK10 effect. As a more specific approach, we next used siRNAs for PAR1 or PAR2, which specifically knocked-down PAR1 and PAR2, respectively (Figure S8a,b). The siRNAs for PAR1 and PAR2 also prevented the anti-inflammatory effect of KLK10, as shown by the monocyte adhesion assay (Figure 6b). When both PAR1 and PAR2 were knockdown together by the siRNAs, no additive effect was observed. To further test the role for PAR1/2 in KLK10-mediated anti-inflammatory effect, we treated PAR1-/-, PAR2-/-, and C57/BL6 wildtype mice with rKLK10 (0.6mg/kg once every 3 days for 1 week). We then performed *en face* immunostaining for VCAM1 in the LC area of the aortic arch. We found that the anti-inflammatory of KLK10 was significantly reduced in PAR1-/- and PAR2-/-mice (Figure S10), confirming the *in vitro* results. Together, these results suggest that both PAR1 and PAR2 are involved in mediating the anti-inflammatory effect of KLK10.

**Figure 6.**
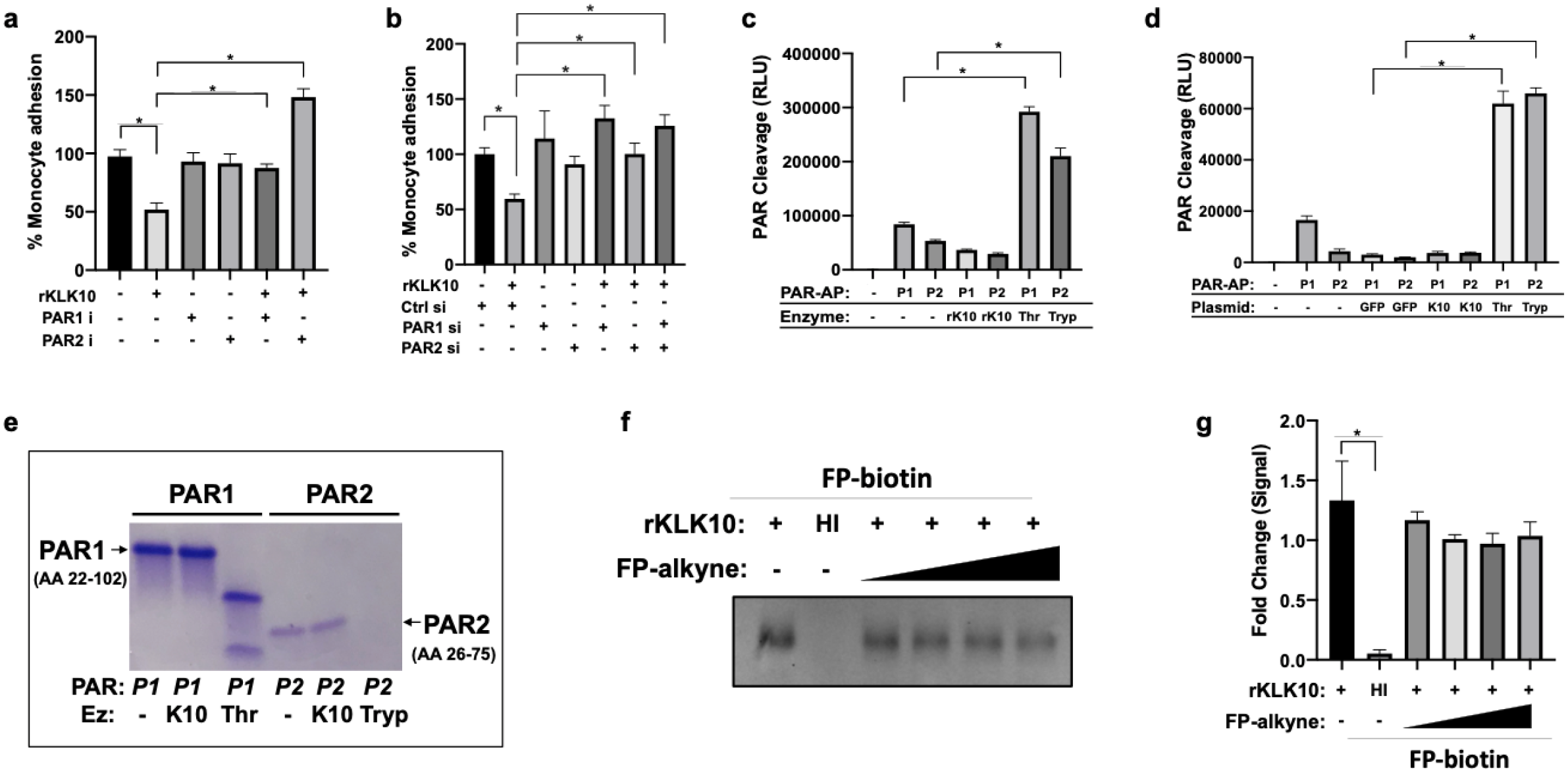
KLK10 inhibits endothelial inflammation in a PAR1/2-dependent manner, but without the direct cleavage. (a) THP1 monocyte adhesion assay was conducted in HAECs subjected to OS (±5 dynes/cm^2^) for 24hrs with rKLK10 and PAR1 (SCH79797, 0.1 μM) or PAR2 inhibitor ( FSLLRY-NH2, 10 μM). Data is normalized to the % of vehicle control. (b) THP1 monocyte adhesion assay was conducted in HAECs subjected to OS (±5 dynes/cm^2^) for 24hrs with rKLK10 and PAR1/2 siRNA or control siRNA. Data is normalized to the % of vehicle control siRNA. (c) PAR cleavage assay in which HAECs were transfected with PAR1-AP or PAR2-AP plasmids and treated with rKLK10 (100 ng/mL), thrombin (5 U/mL) or trypsin (5 U/mL) for 1hr. Conditioned media was then assayed for alkaline phosphatase activity. (d) PAR cleavage assay in which HAECs were co-transfected with PAR1-AP or PAR2-AP plasmids and KLK10 plasmids. Conditioned media was then assayed for alkaline phosphatase activity. (e) Synthetic peptides (100 μM) corresponding to the N-terminal extracellular domains of PAR1 (AA22-102) or PAR2(AA26-75) were incubated with rKLK10 (100 ng/mL), thrombin (5 U/mL) or trypsin (5 U/mL) for 1hr at 37°C in 50 mM Tris, 150 mM NaCl, pH 8.0 and peptide products were analyzed by Tricine SDS-PAGE. (f) KLK10 enzymatic activity was measured by incubating rKLK10 (60 ng) with FP-Biotin (50 μm) and increasing concentrations of FP-alkyne competitive inhibitor (50-500 μm) followed by streptavidin-HRP western blotting. FP-biotin was also incubated with heat-inactivated (HI) rKLK10. (g) Quantification of KLK10 enzymatic activity represented as fold change normalized to FP-Biotin+rKLK10. All data is represented as Mean±SEM, *P≤0.05, One-Way Anova with Bonferroni Correction. N=3-6.

Since PARs are activated by site-specific cleavage of their extracellular N-terminal domain^45, 46^, we tested if KLK10 can also cleave PAR1 and PAR2 to further define the mechanism of action. For this study, we used two independent approaches. First, a secreted alkaline phosphatase (SEAP) reporter fused to the N-terminal domain of either PAR1 or PAR2 (PAR1-AP or PAR2-AP) was expressed in HAECs as reported previously^47–49^. As expected, treatment with thrombin (PAR1 agonist) increased activity of SEAP from the PAR1-AP, indicating the cleavage of the PAR1 N-terminus in response to the canonical agonist (Figure 6c). Similarly, trypsin (PAR2 agonist) also increased the SEAP activity in PAR2-AP expressing HAECs. Unexpectedly, however, treatment with rKLK10 did not induce the SEAP activity, indicating it did not cleave the PAR1 or PAR2. To ensure that rKLK10 was still able to inhibit endothelial inflammation, we concomitantly measured monocyte adhesion under the same condition and observed the expected anti-inflammatory effect (Figure S8c). Next, we tested if KLK10 overexpression using the plasmid vector could cleave PAR1-AP or PAR2-AP in HAECs. Consistent with the rKLK10, KLK10 overexpression also failed to cleave either PAR1-AP or PAR2-obtained using human embryonic kidney cells (not shown).

Next, as an alternative, independent approach, we carried out a peptide cleavage assay using the synthetic N-terminal peptides corresponding to amino acid sequence 22-102 of PAR1 (PAR1^22–102^) and 26-75 of PAR2 (PAR2^26–75^), which contain canonical cleavage-activation sites for known proteinase agonists^45, 46^. Again, as expected, thrombin and trypsin efficiently cleaved the PAR1 and PAR2 peptides, respectively, as demonstrated by the Coomassie staining of Tricine-SDS PAGE gel (Figure 6e). However, rKLK10 failed to expression vector can cleave and activate PAR1 or PAR2 receptors.

We next tested whether rKLK10 has enzymatic activity by incubating rKLK10 with an FP-biotin serine proteinase Activity-Based Probe, followed by streptavidin-HRP western blotting. rKLK10 was labelled with FP-biotin (Figure 6f,e), indicating that KLK10 is indeed an active serine proteinase. Of importance, labeling by the activity-based probe was lost when rKLK10 was heat-inactivated, suggesting its 3D conformation is necessary for its enzymatic activity and the anti-inflammatory effect. Taken together, these results demonstrate that KLK10 inhibits endothelial inflammation in a PAR1/2-dependent manner, but without directly cleaving the PARs.

### KLK10 binds to endothelial proteins HTRA1, VEGFA, and VATB2

Thus far, our results indicated that KLK10 induces anti-inflammatory effects in a PAR1/2-dependent manner without cleaving them. Therefore, we hypothesized that KLK10 binds to another protein which mediates the PAR1/2-dependent anti-inflammatory effect. To test this hypothesis, we carried out an affinity pulldown assay using rKLK10 conjugated to TRICEPS-biotin to identify KLK10 binding proteins in HAECs followed by mass spectrometry^50, 51^. Using glycine conjugated TRICEPS and streptavidin-PE alone as negative controls, we found that rKLK10 coupled to TRICEPS specifically bound to the endothelial membrane as determined by flow cytometry (Figure S11a,b) and immunostaining (Figure S11c,d).

To identify KLK10-binding proteins in HAECs, rKLK10 conjugated to TRICEPS-biotin was pulled down using Streptavidin followed by mass spectrometry. As a positive control, Transferrin conjugated to TRICEPS-biotin was used since it was previously shown to bind the Transferrin receptor TFR1^50, 51^. Mass spectrometry data showed that Transferrin bound to TRF1, as expected (Figure 7a). In contrast, KLK10 bound to HTRA1, VEGFA (Vascular Endothelial Growth Factor A), and VATB2 (ATPase H+ Transporting V1 Subunit B2), as well as KLK10 itself. Of the three KLK10 binding proteins, we selected HTRA1 for further study because of its potential clinical significance and novelty.

**Figure 7.**
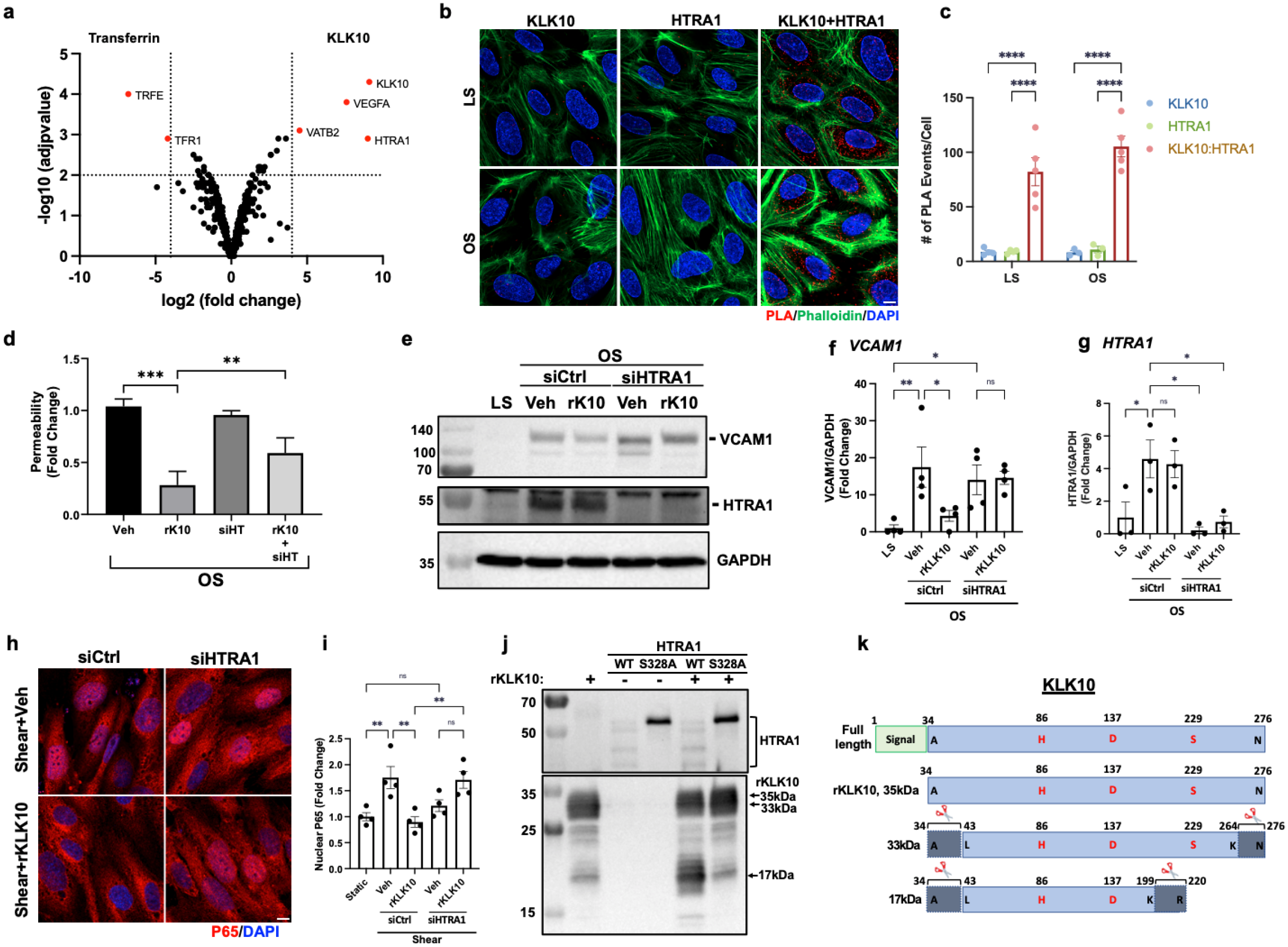
HTRA1 binds and cleaves KLK10 and is necessary for the anti-inflammatory and barrier protective effects of KLK10. (a) Volcano plot depicting proteins that bind to KLK10 compared to the Transferrin (TRFE) control in HAECs using a TRICEPS affinity pulldown method, followed by mass spectrometry. Three KLK10 binding proteins (HTRA1, VEGFA, were VATB2) in addition to KLK10 itself were identified as highlighted in red. (b) The binding of KLK10 and HTRA1 as indicated by red staining was demonstrated by the proximity ligation assay (PLA) using KLK10 and HTRA1 antibodies in HAECs subjected to 48hr of LS or OS. HAECs were then counterstained with Phalloidin D (green) and DAPI (blue). (c) Quantification of the number of PLA interactions per cell is shown, n=5. (d-i) HAECs pre-treated with HTRA1 siRNA (50 nM) were exposed to OS in the presence of rKLK10 (10 ng/mL) for 24hrs, followed by the FITC-avidin permeability assay (d) or Western blots using antibodies to VCAM1, HTRA1, and GAPDH control (e-g). (h,i) HAECs pre-treated with HTRA1 siRNA were exposed to LS (15 dynes/cm^2^) in the presence of rKLK10 (10 ng/mL) for 1hr to induce p65 nuclear localization and immunostained with p65 antibody (red) and DAPI (blue). Quantification of nuclear p65 levels is shown as fold-change over static control, n=4. (j) rKLK10 (500 ng) and rHTRA1 (500 ng; WT or S328A mutant) were incubated for 18hrs, and cleaved proteins were analyzed by western blot using KLK10 and HTRA1 antibodies. (k) Mass spectrometry analysis of tryptic peptides generated from HTRA1 cleavage of rKLK10 revealed the intact rKLK10 (A34-N276) along with KLK10-33kDa (L43-N276) and KLK10-17kDa (L43-K199) cleaved proteins. Bracketed regions indicate HTRA1 cleavage locations. H, D, and S indicate conserved catalytic triad amino acids. All data are represented as Mean±SEM, *P≤0.05 using One-way ANOVA with Bonferroni correction.

### HTRA1 binds KLK10, regulating its anti-inflammatory and barrier protective function

HTRA1 is a serine protease associated with CARASIL (Cerebral Autosomal Recessive Arteriopathy with Subcortical Infracts and Leukoencephalopathy) and AMD (Age-related Macular Degeneration). While CARASIL is associated with loss-of-function mutations of HTRA1, AMD is associated with gain-of-function mutations, suggesting the importance of regulating HTRA1 activity in a tight range in pathobiology^52–55^.

We further validated the binding identified by the mass spectrometry study between KLK10 and HTRA1 by the Proximity Ligation Assay in HAECs exposed to LS or OS (Figure 7b,c). We found that KLK10 and HTRA1 are bound to each other in both LS and OS conditions, with no significant difference between the groups. Next, we tested if HTRA1 has functional importance in the anti-inflammatory and barrier protective function of KLK10 by using the HTRA1 siRNA in HAECs exposed to OS. The siRNA-mediated knockdown of HTRA1 significantly reduced the KLK10’s protective effects on shear-induced permeability (Figure 7d), VCAM1 induction (Figure 7e,f), and p65 NFκB nuclear localization (Figure 7h,i). These results demonstrate that HTRA1 regulates the anti-inflammatory and barrier-protective effects of KLK10. Interestingly, we found that HTRA1 was upregulated by OS in HAECs (Figure 7e,g), indicating HTRA1 itself is shear sensitive. This finding was further validated by qPCR in HAECs (Figure S12a) and by scRNAseq (Figure S13) and scATACseq analyses (Figure S14) in the mouse PCL study^38^. These results show KLK10 and HTRA1 expression are inversely regulated by flow.

### HTRA1 cleaves KLK10 at the N-terminal and C-terminal regions

Since HTRA1 is a serine protease, we tested if HTRA1 cleaves KLK10. To test this hypothesis, we incubated rKLK10 (A34-N276, 35kDa) with recombinant WT or catalytically inactive HTRA1 mutant (S328A), and analyzed the KLK10 cleavage products by western blot and Coomassie staining. Incubation of rKLK10 (35kDa form) with WT rHTRA1 generated two cleaved KLK10 fragments at 33kDa and 17kDa, while incubation with the inactive rHTRA1 mutant continued to have the intact 35kDa KLK10 band (Figure 7j). The mass spectrometry analysis of the tryptic peptides produced from digesting these three bands with trypsin revealed that all three were indeed KLK10 proteins. The tryptic digestion mapping analysis of the mass spectrometry data (Figure 7k and Figure S15) suggested that the 35kDa protein was the intact rKLK10 (A34-N276). In contrast, the 33kDa and 17kDa fragments showed evidence for cleavages at the N-terminus and two different C-termini. The 33kDa form contained L43-K264 (missing A34-R42 at the N-terminus and Y265-N276 at the C-terminus), while the 17kDa form showed L43-K199 (missing A34-R42 at the N-terminus and E200-R220 at the C-terminus). These missing tryptic peptides at both the N- and C-terminal regions of KLK10 may contain potential cleavage sites by HTRA1.

### KLK10 expression is decreased in human coronary arteries with advanced atherosclerotic plaques

We next examined if KLK10 expression is altered in human coronary artery tissue sections with varying degrees of atherosclerotic plaques (n=40 individuals, Table S1). KLK10 and CD31 immunostaining demonstrated that KLK10 expression was significantly reduced at the endothelial layer in arteries with significant plaques (Grade 4-6; Figure 8a,b) than less-diseased arteries (Grades 1-3).

**Figure 8.**
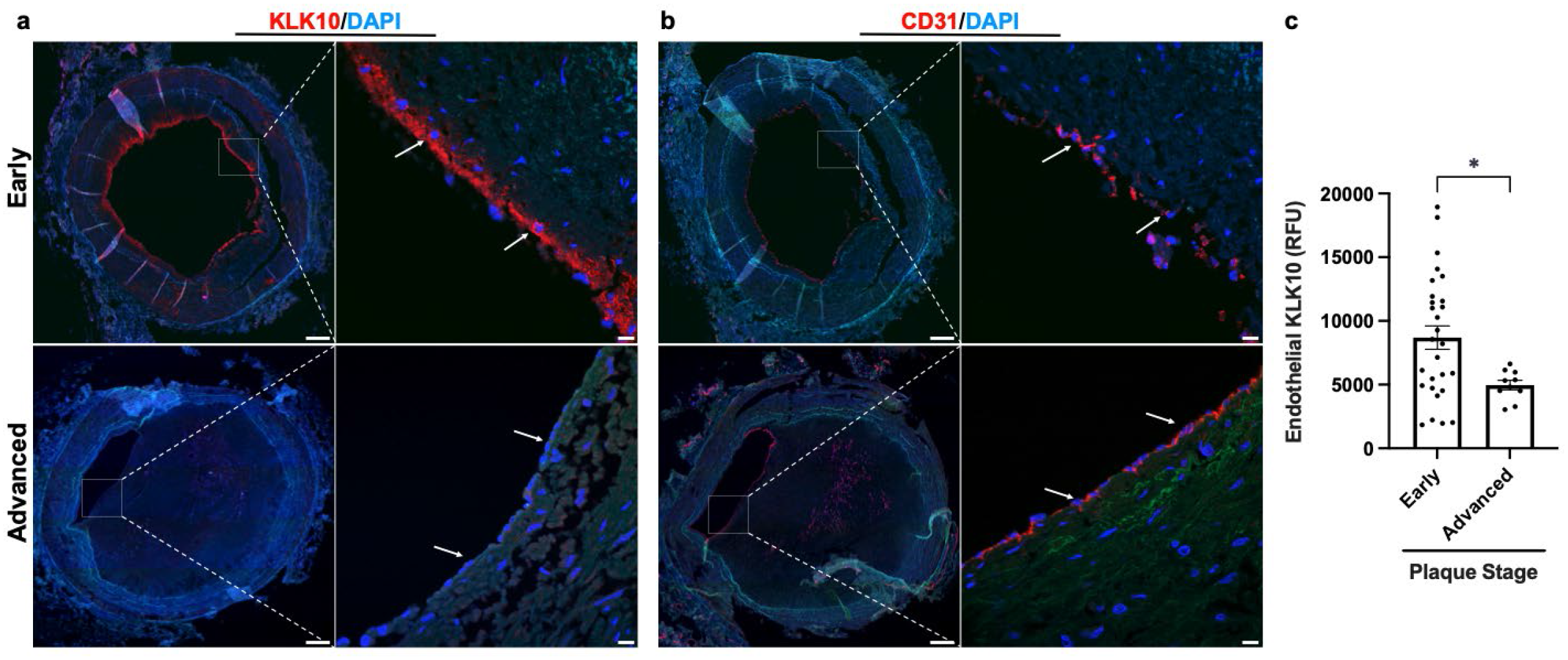
KLK10 expression is decreased in human coronary arteries with advanced atherosclerotic plaques. (a) Human coronary artery sections with varying degrees of atherosclerotic lesions were stained with KLK10 antibody (red) and DAPI (blue). Scale bar low mag=500 μm, scale bar; high mag=50 μm. Arrows indicate endothelial cells. (b) Consecutive arterial sections from the same patients were stained with CD31 antibody (red) and DAPI (blue). (c) Quantification of endothelial KLK10 fluorescent intensity in lower stage plaques (AHA grades 1-3) and advanced stage plaques (AHA grade 4-6). Data are from 40 different patients. Two-tailed t-test. Mean±SEM *P≤0.05.

## Discussion

Here, we describe that *s-flow* promotes, while *d-flow* inhibits, expression and secretion of KLK10 in ECs *in vitro* and *in vivo*. We found for the first time that KLK10 can inhibit endothelial inflammation, endothelial barrier dysfunction, and reduce migration and tube formation, but not apoptosis or proliferation. Importantly, treatment of ECs *in vitro* with rKLK10 or a KLK10 expression plasmid inhibited endothelial inflammation induced by *d-flow* or TNFα. Moreover, treatment with rKLK10 or overexpression of KLK10 by ultrasound-mediated plasmid expression inhibited endothelial inflammation and atherosclerosis development *in vivo*. We found that PAR1/2 mediated the anti-inflammatory effect of KLK10. Unexpectedly, however, but KLK10 did not cleave the PAR1/2, suggesting a non-canonical activation mechanism. Indeed, we found a novel mechanism by which HTRA1 regulates KLK10’s anti-inflammatory and barrier protective functions. Our findings also indicate that KLK10 is likely to be important in human atherosclerotic plaque development. The protective effects of rKLK10 or plasmid-driven KLK10 expression on endothelial inflammation, barrier function, and atherosclerosis suggest its therapeutic potential for atherosclerosis treatment.

Our data revealed that flow regulates the expression of KLK10 and its novel binding partner HTRA1 in an opposite manner, indicating the interaction and balance between the two proteins may play a critical role in KLK10 function. KLK10 expression is downregulated in certain cancers (i.e., breast, prostate, testicular, and lung cancer)^25–29^ but overexpressed in others^30–33^. These findings suggest that abnormal, either too low or too high, levels of KLK10 are associated with various cancer conditions. Similarly, abnormal expression of HTRA1 is implicated in the pathophysiology of other diseases, including CARASIL, AMD, and various cancer^52–55^. Therefore, the balance between HTRA1 and KLK10 may play a critical role in not only endothelial function and atherosclerosis but also in other diseases. Since *s-flow* induced KLK10^high^/HTRA1^low^ while *d-flow* led to KLK10^low^/HTRA1^high^, the KLK10 versus HTRA1 balance could affect KLK10 cleavage status and subsequent activity under different flow conditions. The pathophysiological significance of the KLK10/HTRA1 balance in atherosclerosis and other diseases is an important future question.

We found that HTRA1 cleaves rKLK10 (35kDa form) to generate 33kDa and 17kDa cleavage products. The 33kDa and 17kDa forms displayed potential cleavage regions both at the N-terminus and two different C-termini (Figure 7I,k, S15). Additional studies are required to identify the exact cleavage sites within these regions in ECs and tissues. Interestingly, however, the 17kDa form loses one of the residues involved in the catalytic triad (S229), which may lead to the loss of KLK10 activity. The functional importance of these KLK10 fragments needs to be determined in the future.

Mechanistically, our data using the PAR inhibitors and siRNAs (Figures 6a and 6b) show that PAR1/2 mediate the anti-inflammatory effect of KLK10 in ECs. Since PAR1/2 can be activated by KLKs 5, 6, and 14^43, 44^, it came as a surprise that KLK10 was not able to cleave and activate the PARs directly in our study using several independent approaches (Figure 6). This was demonstrated by SEAP-PAR1/2 or N-Luciferase-PARs 1/2 cleavage assay in response to either rKLK10 or the KLK10 expression vector, as well as the cleavage assays using the PAR1/2 peptides representing the canonical cleavage sites. Nonetheless, the activity-based probe analysis showed that KLK10 was indeed catalytically active. Of note, KLK10 lost its enzymatic activity and the anti-inflammatory effect upon heat-denaturation, indicating the importance of the 3D conformation for its enzyme activity and biological activity.

Taken together, our results show that KLK10’s anti-inflammatory effect in ECs does not require the direct cleavage of PAR1/2. Instead, HTRA1 binds, cleaves, and possibly activates KLK10, regulating its anti-inflammatory response in a non-canonical PAR1/2-dependent manner (Figure 9). The interaction of KLK10 and HTRA1 leads to inhibition of the NFκB activation and endothelial inflammation through the non-canonical PAR1/2-dependent manner, and atherosclerosis (Figure 9). The mechanism by which KLK10 and HTRA1 interaction regulates the PAR signaling pathway is an important unanswered question.

**Figure 9.**
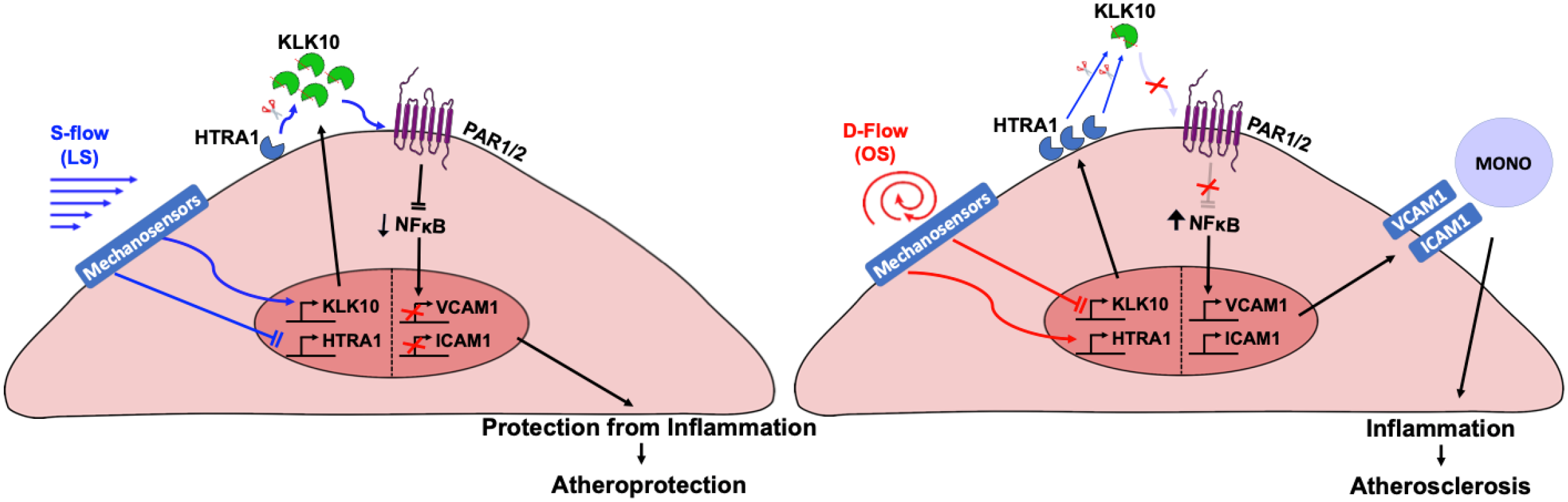
Working hypothesis: Flow-sensitive KLK10 inhibits inflammation and atherosclerosis in a HTRA1- and PAR1/2-dependent mechanism. KLK10 is upregulated by *s-flow* and downregulated by *d-flow*. HTRA1 is also a secreted serine protease with inverse regulation by flow relative to KLK10. Under *s-flow* conditions when KLK10 is expression is high and HTRA1 expression is low, HTRA1 may cleave and activates KLK10, which in turn activates non-canonical PAR1/2 pathway to inhibit NFκB-mediated expression of VCAM1 and ICAM1, thereby decreasing monocyte adhesion and atherosclerosis. Under *d-flow* conditions when KLK10 is expression is low and HTRA1 expression is high, HTRA1 cleaves KLK10 to prevent its anti-inflammatory and anti-atherogenic pathway.

In summary, we show that KLK10 is a flow-sensitive protein that is upregulated by *s-flow* and downregulated by *d-flow* in ECs. Our results also demonstrate that KLK10, working with HTRA1, is a key mediator of the anti-inflammatory and anti-atherogenic effects of *s-flow* in a PAR1/2-dependent manner. The current study also raises numerous new and unexpected questions regarding the role and mechanisms of KLK10 and HTRA1 in vascular pathophysiology. KLK10 and HTRA1 may serve as potential anti-atherogenic therapeutic targets.

## Materials and Methods

### Mouse studies

All animal studies were performed with male C57BL/6 or *ApoE*^-/-^ mice (Jackson Laboratory), were approved by Institutional Animal Care and Use Committee by Emory University, and were performed in accordance with the established guidelines and regulations consistent with federal assurance. All studies using mice were carried out with male mice at 6-10 weeks to reduce the sex-dependent variables. For partial carotid ligation studies, mice at 10 weeks were anesthetized and 3 of 4 caudal branches of LCA (left external carotid, internal carotid, and occipital artery) were ligated with 6-0 silk suture, but the superior thyroid artery was left intact. Development of *d-flow* with characteristic low and oscillating shear stress in each mouse was determined by ultrasound measurements as we described^14^. Following the partial ligation, mice were either continued to be fed chow-diet for 2 days or high-fat diet for atherosclerosis studies for 3 weeks as specified in each study.

Endothelial-enriched RNA was prepared from the LCA and the contralateral RCA control following 48hrs after the partial ligation as we described previously^14^. For *en face* immunostaining, mice were euthanized under CO_2_ and the aortas were pressure-fixed with 10% formalin saline^14^. The aortas were carefully cleaned *in situ*, and the aortic arches and thoracic aortas were dissected, fixed in ice-cold acetone for 5 minutes, permeabilized using 0.1% Triton-X100 in PBS for 15 minutes, blocked for 2hrs with 10% donkey serum, and incubated with anti-KLK10 (BiossUSA bs-2531R, 1:100) and anti-VCAM1 (Abcam ab134047, 1:100) primary antibodies overnight at 4°C followed by Alexa Fluor-647 secondary antibodies (ThermoFisher Scientific, 1:500) for 2hrs at room temperature. The aortas were opened and the lesser curvature (LC) and greater curvature (GC) of each arch were separated. The aortas were then mounted on glass slides with VectaShield that contained DAPI (Vector Laboratories). *En face* images were collected as a Z-stack with a Zeiss LSM 800 confocal microscope. For mouse frozen section staining studies, fresh mouse aortas were placed in Tissue-Tek OCT compound, snap-frozen in liquid nitrogen, and sectioned at 7 μm as we described^56^.

### Immunohistochemical staining of sections from human coronaries

For human coronaries arteries, 2 mm cross sections of the left anterior descending arteries were obtained from de-identified human hearts not suitable for cardiac transplantation donated to LifeLink of Georgia. The de-identified donor information is shown in Supplementary Table 1. Tissues were fixed in 10% neutral buffered formalin overnight, embedded in paraffin, and 7 μm sections were taken, and stained as we described^37, 57^. Sections were deparaffinized and antigen retrieval was performed as described previously^37, 57^. Sections were then permeabilized using 0.1% Triton-X100 in PBS for 15 minutes, blocked for 2hrs with 10% goat serum, and incubated with anti-KLK10 (BiossUSA bs-2531R, 1:100) primary antibody overnight at 4°C followed by Alexa Fluor-647 (ThermoFisher Scientific, 1:500) secondary antibody for 2hrs at room temperature. Nuclei were counter-stained with DAPI (Vector Laboratories, Burlingame, Calif). Hematoxylin and Eosin staining (American Mastertech) and plaque area quantification using ImageJ software (NIH) were done as we described^37, 57^. All confocal images were taken with a Zeiss (Jena, Germany) LSM800 confocal microscope.

### Cell culture and *in vitro* shear stress study

HAECs were obtained from Lonza and maintained in EGM2 medium (Lonza) supplemented with 10% fetal bovine serum (Hyclone), 1% bovine brain extract, 10 mM L-glutamine, 1 μg/mL hydrocortisone hemisuccinate, 50 μg/mL ascorbic acid, 5 ng/mL EGF, 5 ng/mL VEGF, 5 ng/mL FGF, and 15 ng/mL IGF-1 as we described^56^. HUVECs were purchased from BD Biosciences, cultured in M199 media (Cellgro) supplemented with 20% fetal bovine serum (Hyclone), 1% bovine brain extract, 10 mM L-glutamine, and 0.75 U/mL heparin sulfate as we described^58^. All ECs were grown at 5% CO2 and 37°C and used between passages *5* and *9*. THP-1 monocytes were obtained from ATCC and maintained in RPMI-1640 medium supplemented with 10% FBS and 0.05 mM 2-mercaptoethanol at 5% CO2 and 37°C as we described^58^. For flow experiments, confluent HAECs or HUVECs were exposed to steady unidirectional laminar shear stress (LS, 15 dyn/cm^2^) or bidirectional oscillatory shear stress (OS, ±5 dyn/cm^2^ at 1 Hz), mimicking *s-flow* and *d-flow* conditions, respectively, using the cone-and-plate viscometer for 24hrs experiments, as we reported^36, 37^.

### rKLK10 and KLK10 Plasmids

Initially, human rKLK10 (Ala34-Asn276 with a 6x N-terminal His tag) produced in E.coli (Ray Biotech, 230-00040-10) were used. Additional studies using human rKLK10 produced in the mammalian CHO-K1 cells validated the initial results. Most studies were carried out using human rKLK10 produced in CHO-K1 cells using a full-length expression vector (pcDNA 3.4, Met1-Asn276). rKLK10 with a 6x C-terminal His tag was affinity purified using HisPur Ni-NTA Resin (Thermo Scientific) per the manufacturer’s instruction (Figure S16) using the conditioned medium. Amino acid sequencing analysis of the purified rKLK10 by mass spectrometry showed that our rKLK10 preparation was a mature form expressing Ala34-Asn276 (data not shown).

### Overexpression or knockdown experiments *in vitro*

Cells were transiently transfected with a human KLK10-encoding plasmid (pCMV6-KLK10-Myc-DDK; Origene RC201139) at 0.1-1 μg/mL or as a control a GFP plasmid (PmaxGFP, Lonza, Cat. No. D-00059) using Lipofectamine 3000 (Invitrogen, Cat. No. L3000008) as we described^56^. Alternatively, cells were transfected with KLK10 siRNA (25 nM; Dharmacon; J-005907-08), PAR1 (50 nM; Dharmacon; L-005094-00-0005), PAR2 (50 nM; Dharmacon; L005095-00-0005), HTRA1 (50 nm; Dharmacon L-006009-00-0010) or control non-targeting siRNA (25 or 50 nM; Dharmacon; Cat. No. D-001810-10-20) using Oligofectamine (Invitrogen, Cat. No. 12252011) as we described^56^.

### Quantitative Real-Time Polymerase Chain Reaction (qPCR)

Total RNAs were isolated using RNeasy Mini Kit (Qiagen 74106) and reverse transcribed to cDNA using High-Capacity cDNA Reverse Transcription Kit (Applied Biosystems 4368814). qPCR was performed for genes of interests using VeriQuest Fast SYBR QPCR Master Mix (Affymetrix 75690) with custom designed primers (Supplementary Table 2) using 18S as house-keeping control as we previously described^56^.

### KLK10 ELISAs

KLK10 secreted into the conditioned cell culture media from HAECs exposed to shear stress was measured by using a human KLK10 ELISA kit (MyBioSource, MBS009286). KLK10 in mouse plasma was measured by using a mouse KLK10 ELISA kit (Antibodies-online, ABIN628061).

### Endothelial Functional Assays

Endothelial migration was measured by the endothelial scratch assay, as we described^59^. Briefly, HUVECs were treated with rKLK10 at increasing doses overnight and cell monolayers were scratched with a 200-μL pipette tip. The monolayer was washed once, and the medium was replaced with 2% serum media. After 6hrs, the number of cells migrated into the scratch area were quantified microscopically using NIH ImageJ.

Endothelial apoptosis was determined using the TUNEL apoptosis assay, as we described^60^. Briefly, HUVECs were treated with rKLK10 at increasing doses overnight and the cells were fixed using 4% paraformaldehyde for 15 minutes and permeabilized with 0.1% Triton X-100 for 15 minutes. TUNEL staining was then performed using a commercially available kit (Roche, 12156792910) and the number of TUNEL-positive cells were counted using NIH ImageJ.

Endothelial proliferation was determined using Ki67 immunohistochemistry, as we described^61^. Briefly, HUVECs were treated with rKLK10 at increasing doses overnight and the cells were washed twice with PBS, fixed using 4% paraformaldehyde for 15 minutes, and permeabilized with 0.1% Triton X-100 for 15 minutes. After blocking with 10% Goat Serum for 2hrs at RT, cells were incubated overnight at 4°C with rabbit anti-Ki67 primary antibody (Abcam ab15580, 1:100). The following day, cells were washed three times with PBS, incubated for 2hrs at RT protected from light with Alexa-fluor 647-labeled goat anti rabbit IgG (1:500 dilution), and counterstained with DAPI. The number of Ki67 positive cells were counted using NIH ImageJ.

Endothelial tube formation was measured using a Matrigel tube formation assay, as we described^59^. Briefly, HUVECs were seeded in a growth factor reduced Matrigel (BD Bioscience) coated 96-well plate and incubated with rKLK10 (100 ng/mL) for 6hrs at 37°C. Tubule formation was quantified microscopically by measuring tubule length using NIH ImageJ.

Endothelial permeability was determined by FITC-avidin binding to biotinylated gel, as previously described^39^. Briefly, HAECs were seeded on biotinylated-gelatin and treated with rKLK10 overnight followed by thrombin (5 U/mL) for 4hrs or OS for 24hrs as described above. Following the completion of the experiments, FITC-avidin was added to the cells and fluorescent intensity was measured using NIH ImageJ.

Monocyte adhesion to ECs was determined using THP-1 monocytes (ATCC TIB-202) as we described^56^. In brief, THP-1 cells (1.5×10^5^ cells/mL) were labeled with a fluorescent dye 2’,7’-bis(carboxyethyl)-5 (6)-carboxyfluorescein-AM (Thermo Fisher Scientific B1150; 1 mg/mL) in serum-free RPMI medium (Thermo Fisher Scientific 11875093) for 45 minutes at 37°C. After exposure to flow or other experimental treatments, the ECs were washed in RPMI medium before adding 2’,7’-bis(carboxyethyl)-5 (6)-carboxyfluorescein-AM– loaded THP-1 cells. After a 30-minute incubation at 37°C under no-flow conditions, unbound monocytes were removed by washing the endothelial dishes 5× with HBSS and cells with bound monocytes were fixed with 4% paraformaldehyde for 10 minutes. Bound monocytes were quantified by counting the number of labeled cells at the endothelium under a fluorescent microscope.

### Preparation of whole-cell lysate and immunoblotting

After treatment, cells were washed 3× with ice-cold HBSS and lysed with RIPA buffer containing protease inhibitors (Boston Bioproducts BP-421)^56^. The protein content of each sample was determined by Pierce BCA protein assay. Aliquots of cell lysate were resolved on 10% to 12% SDS-polyacrylamide gels and subsequently transferred to a polyvinylidene difluoride membrane (Millipore). The membrane was incubated with the following primary antibodies: anti-KLK10 (BiossUSA bs-2531R, 1:1000) anti-GAPDH (Abcam ab23565, 1:2000), anti-β-actin (Sigma-Aldrich A5316, 1:2000), anti-VCAM1 (Abcam ab134047, 1:1000), anti-ICAM1 (Abcam ab53013, 1:1000), anti-phospho-NFκB (Cell Signaling #3033, 1:1000), and anti-HTRA1 (R&D Systems MAB2916, 1:1000) ( overnight at 4°C in 5% milk in TBST at the concentration recommended by the manufacturer, followed by secondary antibody addition for 1h at RT in 5% milk in TBST. Protein expression was detected by a chemiluminescence method^56^.

### PAR cleavage assays

Synthetic peptides corresponding to the extracellular domain of PAR1 (AA Ala22-Thr102) and PAR2 (AA Ile26-Thr75) were assembled by automated Fmoc/tBu-solid-phase synthesis (model CS336X; CSBio) followed by cleavage in trifluoroacetic acid (TFA)/phenol/thioanisole/ethanedithiol/water (10:0.75:0.5:0.25:0.5, w/w; 25 °C, 90 minutes) and precipitation with diethyl ether. The crude peptides were purified by reversed-phase high-pressure liquid chromatography and were obtained in the form of their TFA salts. Their masses as well as masses of their theoretical Lys-C protease (Promega) fragments, obtained upon digestion, were confirmed by electrospray-ionization mass spectrometry (maXis ESI-TOF; Bruker). PAR1 and PAR2 peptides (100 μM) were then incubated with rKLK10 (100 ng/mL), thrombin (5 U/mL), or trypsin (5 U/mL) for 30 minutes at 37°C and analyzed by Tricine-SDS-PAGE followed by Coomassie stain^62^.

PAR1/2-Alkaline Phosphatase (AP) constructs made as previously described^47–49^ were transfected into HAECs (1 μg/mL) for 24hrs using Lipofectamine 3000. Cells were treated with rKLK10 (100 ng/mL), thrombin (5 U/mL), or trypsin (5 U/mL) for 30 minutes. Conditioned media were then collected and analyzed for secreted alkaline phosphatase activity (SEAP) using a commercial kit (T1015, Invitrogen) and a microplate reader^47–49^.

### rKLK10 treatment and KLK10 overexpression in C57BL/6 and *ApoE*^-/-^ mice

Two independent methods were used, rKLK10 and KLK10 plasmid, to treat mice with KLK10. Treatment with rKLK10 was first performed in C57BL/6 mice by administering rKLK10 (0.006-0.6 mg/kg) by tail-vein once every two days and sacrificed on day five. At the completion of the study, mice were euthanized by CO_2_ inhalation and *en face* preparation of the aorta was performed as we described^56^. Alternatively, *ApoE*^-/-^ on a high-fat diet containing 1.25% cholesterol, 15% fat and 0.5% cholic acid were given the partial carotid ligation surgery and rKLK10 or vehicle was administered by tail-vein once every three days for three weeks as we described^56^. Following the completion of the study, mice were euthanized by CO_2_ inhalation and the aortas were excised, imaged, and sectioned for IHC^56^.

KLK10 plasmid overexpression was performed using ultrasound-mediated sonoporation method of gene therapy as reported^40–42^. Briefly, perfluoropropane microbubbles encapsulated by DSPC and DSPE-PEG2000 (9:1 molar ratio) were made using the shaking method as previously described^40–42^. KLK10 plasmid expressing secreted KLK10 and luciferase (pCMV-Igκ-KLK10-T2A-Luc) from GENEWIZ or luciferase plasmid (pCMV-Luc) from Invitrogen (50 μg each) was then mixed with the microbubbles (5×10^5^) and saline to reach 20 μl total volume. Following partial carotid ligation, *ApoE*^-/-^ mice were intramuscular injected to the hind-limbs with the plasmid-microbubble solution. The injected areas of the hind-legs were then exposed to ultrasound (0.35 W/cm^2^) for 1 minute, and repeated 10 days later. At the completion of the study 3 weeks after the partial ligation and on high-fat diet, mice were anesthetized, administered with luciferin (IP; 3.75 mg) and imaged for bioluminescence on a Bruker *In Vivo* Xtreme X-ray Imaging System. Mice were then euthanized by CO_2_ inhalation and the aortas were excised, imaged, and sectioned for staining as described above.

### KLK10 affinity pulldown using TRICEPS

Affinity pulldown using TRICEPS-biotin was performed as described previously^51^. Briefly, HAECs were grown to confluency on a 15 cm^2^ dish and were oxidized using 1.5 mM sodium metaperiodate (NaIO4) for 15min at 4°C. rKLK10, glycine, or Transferrin (300 μg each) was coupled to 150 μg of TRICEPS v.3.0 (DualSystems Biotech) for 1.5hr in 25 mM HEPES pH 8.2 at room temp and added to the HAECs for 1.5hr at 4°C. For flow cytometry and immunostaining experiments, cells were fixed with 4% paraformaldehyde and incubated with Streptavidin-PE for 1hr followed by flow cytometry or confocal microscopy. For the identification of KLK10 binding proteins, cells were scraped and lysed, followed by affinity pulldown and submitted to DualSystems Biotech for LC-MS/MS tandem mass spectrometry analysis as previously described^51^.

### Proximity Ligation Assay

Duolink In Situ Red Starter Kit Mouse/Rabbit Proximity Ligation Assay was purchased from Millipore Sigma (DUO92101) and was performed according to the manufactures protocol using KLK10 and HTRA1 antibodies described above.

### rKLK10:HTRA1 cleavage assay and mass spectrometry analysis of KLK10 cleavage products

rKLK10 (500 ng, described above) was incubated with wild-type human recombinant (rHTRA1) (500 ng, Origene TP322362) or kinase-inactive rHTRA1 containing an S328A mutation (500 ng Origene TP700208) for 18hrs at 37°. Cleavage products were reduced with dithiothreitol (5 mM) for 30min at room temp followed by alkylation with iodoacetamide (25 mM) for 30min at room temp in the dark and run on a 12% SDS-PAGE gel followed by Coomassie G-250 staining. Gel pieces corresponding to 35, 33, and 17kDa KLK10 proteins were excised from Coomassie-stained gels for tryptic digestion and mass spectrometry.

### In-gel digestion

Gel pieces were cut into smaller pieces for efficient destaining and digestion. The pieces were covered with 50 mM ammonium bicarbonate/acetonitrile (1:1, v/v) and were incubated with occasional vortexing for 5 min, after which the solution was discarded. This destaining step was repeated depending on the staining intensity. Subsequently, neat acetonitrile was added and the pieces were incubated with occasional vortexing for 10 min or until gel pieces became white and dry. The acetonitrile was discarded and the gel pieces were dried under vacuum. For digestion, the gel pieces were covered with trypsin solution (10 ng/µL) in 50 mM ammonium bicarbonate and were incubated overnight at 37 °C. Next day, the supernatant containing the tryptic peptides were saved and the gel pieces were used for further extraction of peptides. Gel pieces were completely covered with the extraction buffer containing (1:1, v/v) 5% formic acid/50% acetonitrile and were incubated with occasional vortexing for 10 min. The extraction was repeated one more time followed by incubation with neat acetonitrile. The extracted peptides were dried under vacuum.

### Mass Spectrometry

Dried peptides were resuspended in 10 μL of loading buffer (0.1% trifluoroacetic acid, TFA) and 2 μL was loaded onto a self-packed 25 cm (100 μm internal diameter packed with 1.7 μm Water’s CSH beads) and was separated using Dionex 3000 RSLCnano liquid chromatography system. The liquid chromatography gradient started at 1% buffer B (buffer A: 0.1% formic acid in water, buffer B: 0.1 % formic acid in acetonitrile) and ramps to 5% in 0.1 minute. This was followed by a 25 min linear gradient to 40% B and finally a 5 minute 99% buffer B flush. The mass spectrometry data were acquired on an Orbitrap Fusion Tribrid Mass Spectrometer (ThermoFisher Scientific). The spectrometer was operated in data dependent mode in top speed with a cycle time of 3 seconds. Survey scans were collected in the Orbitrap with a 60,000 resolution, 400 to 1600 m/z range, 400,000 automatic gain control (AGC), 50 ms max injection time and RF lens at 60%. Higher energy collision dissociation (HCD) tandem mass spectra were collected in the ion trap with a collision energy of 35%, an isolation width of 1.6 m/z, AGC target of 10000, and a max injection time of 35 ms. Dynamic exclusion was set to 30 seconds with a 10 ppm mass tolerance window.

### Protein identification

Mass spectrometry data was analyzed according to a published protocol^63^. Spectra were searched using Proteome Discoverer 2.1 against human Uniprot (Swiss-Prot only) database (20,381 target sequences). Searching parameters included fully tryptic restriction and a precursor mass tolerance (± 20 ppm). Methionine oxidation (+15.99492 Da), asaparagine and glutamine deamidation (+0.98402 Da) and protein N-terminal acetylation (+42.03670) were variable modifications (up to 3 allowed per peptide); cysteine was assigned a fixed carbamidomethyl modification (+57.021465 Da). Percolator was used to filter the peptide spectrum matches to a false discovery rate of 1%.

### Serum lipid analysis

Serum lipid analysis was performed at the Cardiovascular Specialty Laboratories (Atlanta, GA) using a Beckman CX7 biochemical analyzer for total cholesterol, triglycerides, HDL and LDL as we reported^56^.

### Statistical analyses

Statistical analyses were performed using GraphPad Prism software. All of the n numbers represent biological replicates. Error bars depict the standard error of means (SEM). Initially, the data sets were analyzed for normality using the Shapiro-Wilk test (*P*<0.05) and equal variance using the *F* test (*P*>0.05). Data that followed a normal distribution and possessed equal variance were analyzed using 2-tailed Student *t* test or 1-way ANOVA, where appropriate with Bonferroni post hoc test as needed. In the case where the data showed unequal variances, an unpaired *t* test with Welch correction was performed or Brown-Forsythe and Welch ANOVA for multiple comparisons. In the case where the data failed the Shapiro-Wilk test (*P*>0.05), a nonparametric Mann-Whitney *U* test was conducted for pairwise comparisons or the Kruskal-Wallis for multiple groups was performed. For animal experiments, we carried out a priori power analysis (G*Power3 analysis at d=2.0, α=0.05, power=0.95, one-tail *t*-test) based on our previous experience using the PCL model to determine the number of animals needed to achieve statistically significant results. Outliers were determined and excluded by calculating interquartile range (IQR) and multiplying 1.5 x IQR.

## Acknowledgment

This work was supported by funding from the National Institutes of Health grants HL119798 and HL095070 to HJ. DW was supported by the NIH F31 HL145974 grant. HJ was also supported by John and Jan Portman Professorship and Wallace H. Coulter Distinguished Faculty Professorship. LOM was supported by the NIH grants HL104165 and HL142975. EWT and LZ. thank Cancer Research UK for support (grant C24523/A25192). JP was supported by the Centers for Disease Control and Prevention (CDC) LaSSI 201706. Excellent technical support of Priyanka Shakamuri and Jaclyn Weinberg at the CDC in synthesizing PAR1 and PAR2 peptides is acknowledged.

---

**Figure S1.**
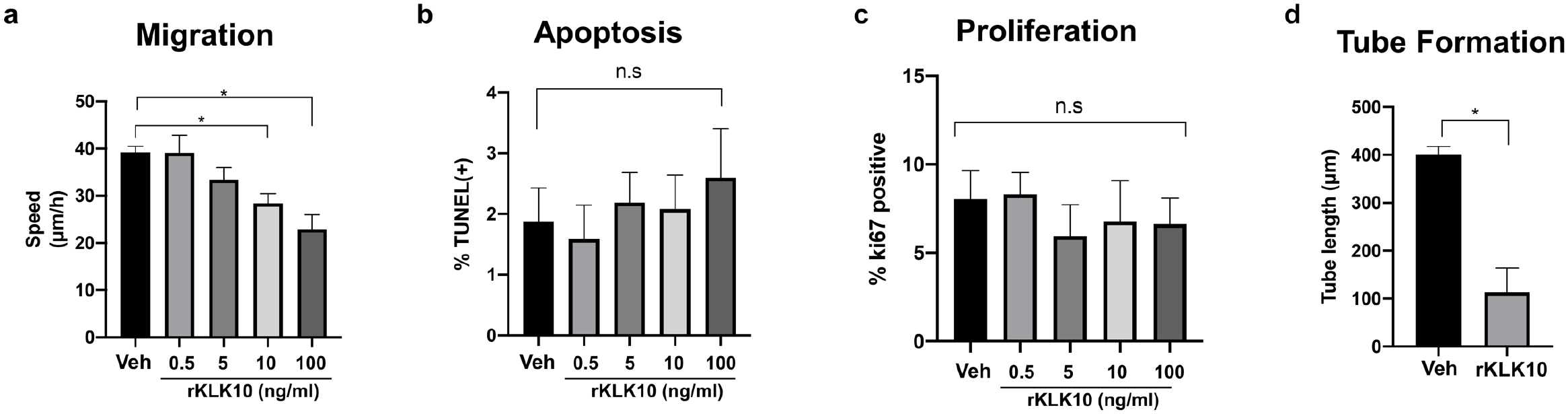
KLK10 inhibits endothelial migration and tube formation, but not apoptosis or proliferation. Human umbilical vein endothelial cells (HUVECs) were treated with rKLK10 from 0.5-100 ng/mL and (a) the scratch assay was performed to measure the rate at which endothelial cells migrated across the scratch; (b) apoptosis was assessed by TUNEL staining; (c) proliferation was assayed by ki67 imunnostaining. (d) HUEVCs were grown on Matrigel and treated with rKLK10 at 100 ng/mL or vehicle and tube length was measured in ImageJ. One-way ANOVA with Bonferroni correction for multiple comparisons where appropriate (a-c) or paired two-tailed t-test (d). Mean±SEM. n=4-6 *P≤0.05.

**Figure S2.**
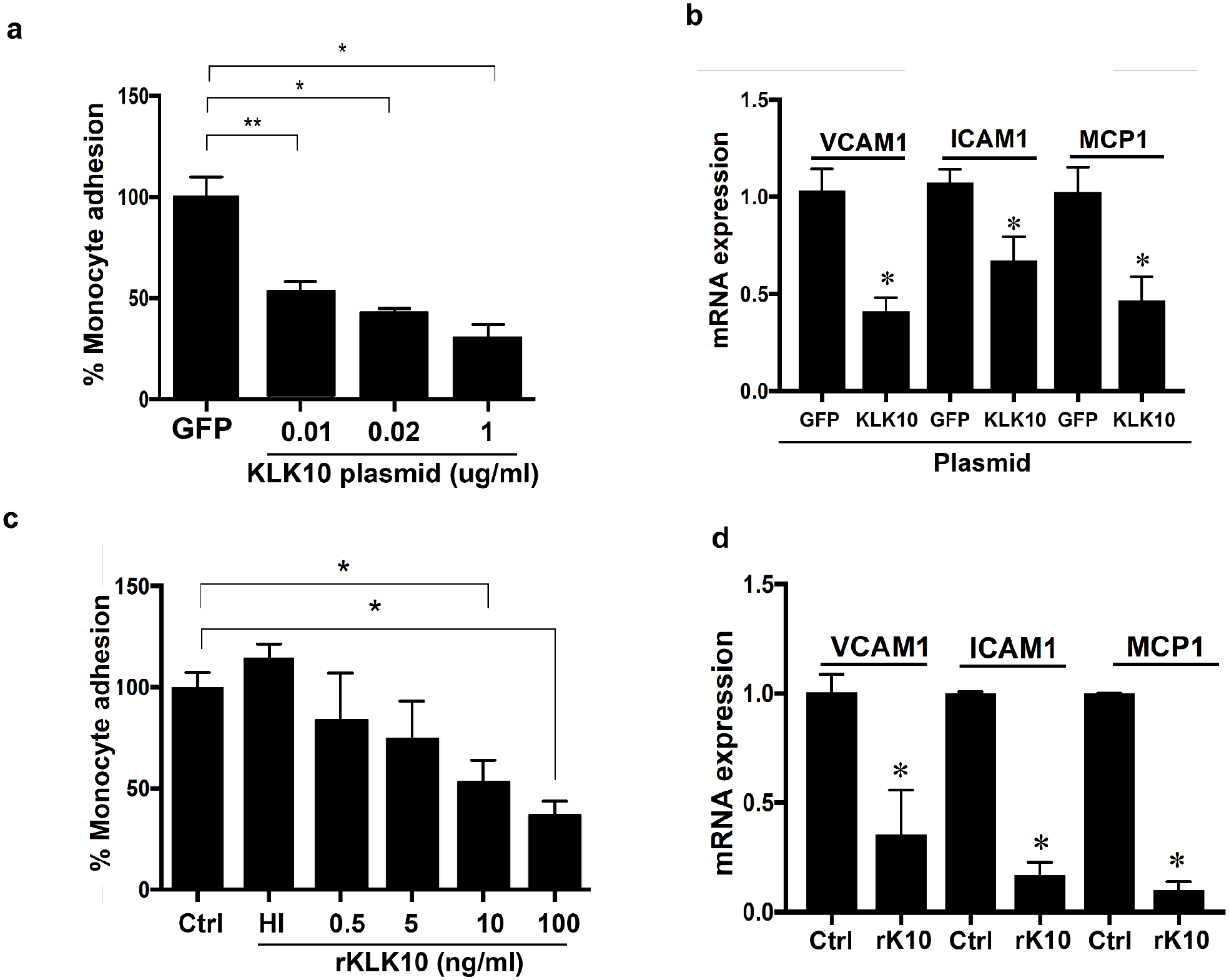
KLK10 reduces inflammation in endothelial cell. (a) Human aortic endothelial cells (HAECs) were transfected with KLK10 plasmid ranging from 0.01-1 μg/mL for 24h and the THP1 monocyte adhesion assay was performed. (b) HAECs were transfected with 1 μg/mL KLK10 plasmid for 24h and qPCR was performed to assess mRNA expression of VCAM1, ICAM1, and MCP1. (c) HAECs were treated with 0.5 to 100 ng/mL rKLK10 and monocyte adhesion assay was performed. (d) HAECs were treated with 100 ng/mL rKLK10 for 24h and qPCR was performed to assess mRNA expression of VCAM1, ICAM1, and MCP1. One-way ANOVA with Bonferroni correction for multiple comparisons (a,c) or two-way ANOVA with Bonferroni correction for multiple comparisons (b,d). Mean±SEM. n=6. *P≤0.05.

**Figure S3.**
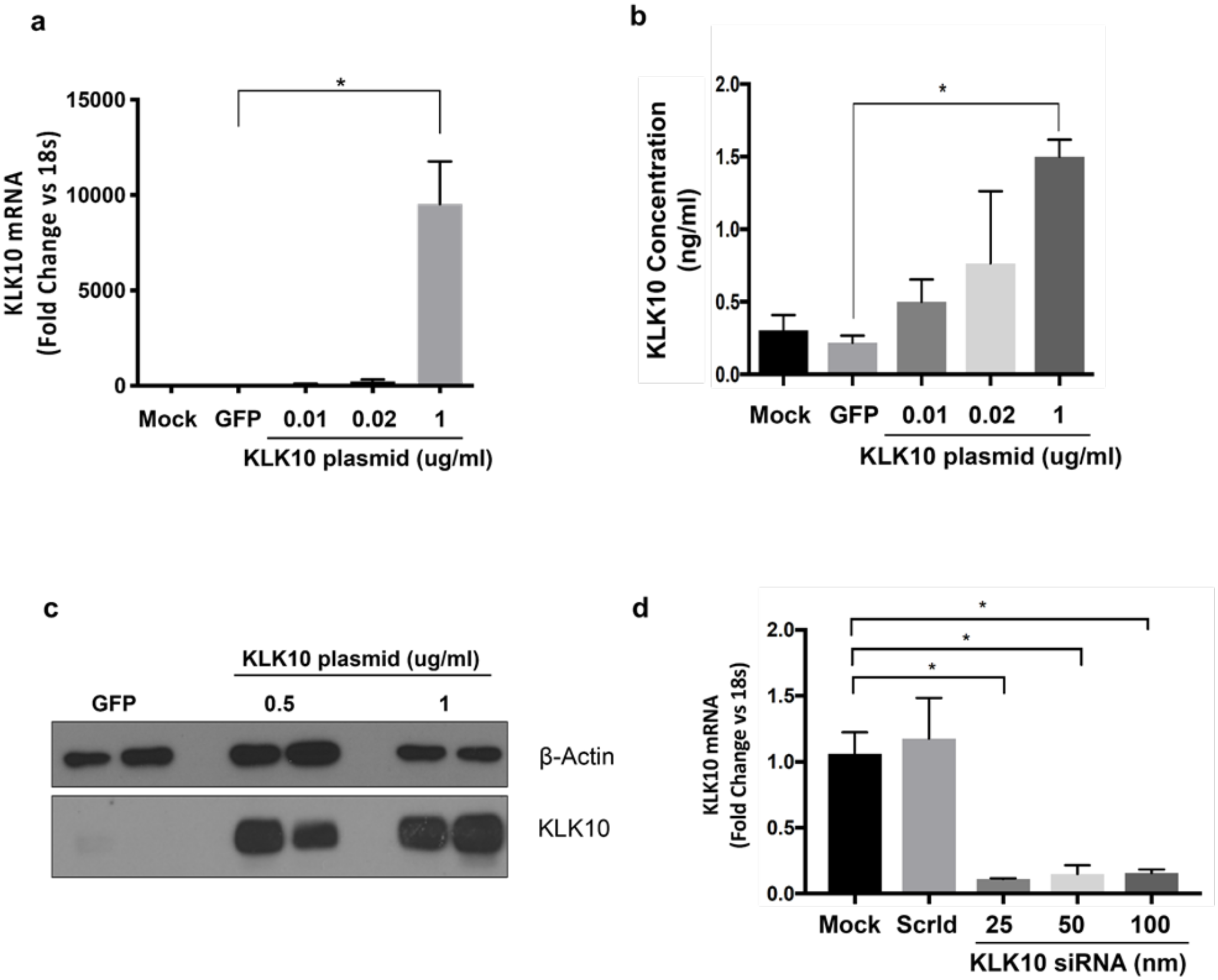
KLK10 plasmid and KLK10 siRNA overexpress and knockdown KLK10, respectively. (a) Human Aortic Endothelial Cells (HAECs) were transfected with 0.02-1 μg/mL KLK10 plasmid or 1 μg/mL GFP plasmid and KLK10 mRNA expression was measured by qPCR. (b) HAECs were transfected with 0.02-1 μg/mL KLK10 plasmid or 1 μg/mL GFP plasmid and KLK10 secretion into the media was measured by ELISA. (c) HAECs were transfected with 500 ng/mL or 1 μg/mL KLK10 plasmid and KLK10 protein expression was measured by western blot, using B-actin as an internal control. (d) HAECs were transfected with 25-100 nM KLK10 siRNA or 100nM scrambled siRNA and KLK10 mRNA expression was measured by qPCR. n=4. Two tailed two-way ANOVA with Bonferroni correction. One-way ANOVA with Bonferroni correction for multiple comparisons where appropriate. Mean±SEM. *P≤0.05.

**Figure S4.**
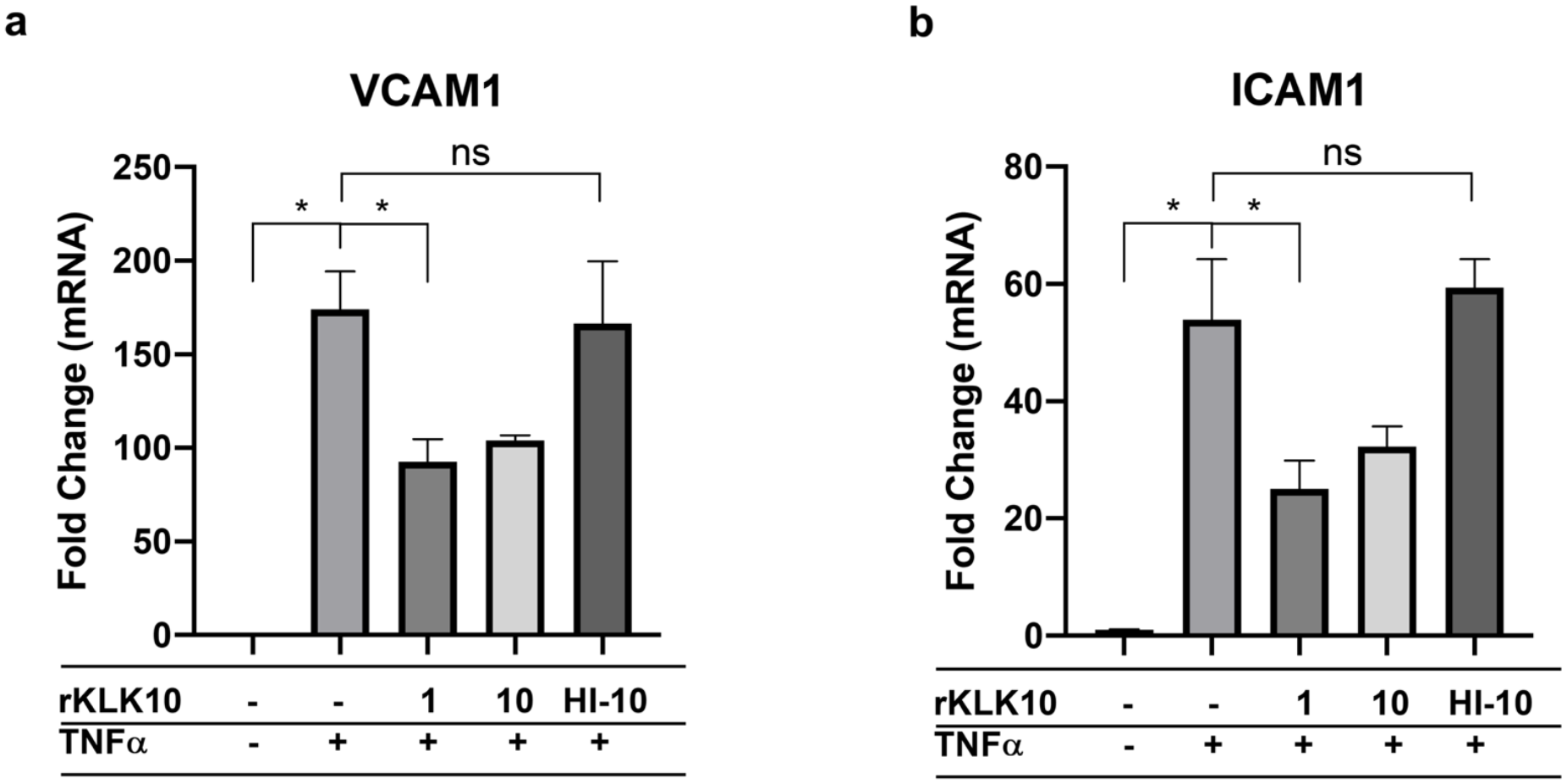
Heat-inactivation of rKLK10 prevents its anti-inflammatory effects on VCAM1 and ICAM1 expression. (a,b) HAECs were treated with TNFα (5 ng/mL) for 4h followed by 1 ng/mL, 10 ng/mL, or 10 ng/mL heat-inactivated (HI) rKLK10 overnight and mRNA expression of (a) VCAM1 and (b) ICAM1 was measured by qPCR. n=6. One-way ANOVA with Bonferroni correction for multiple comparisons. Mean±SEM. *P≤0.05.

**Figure S5.**
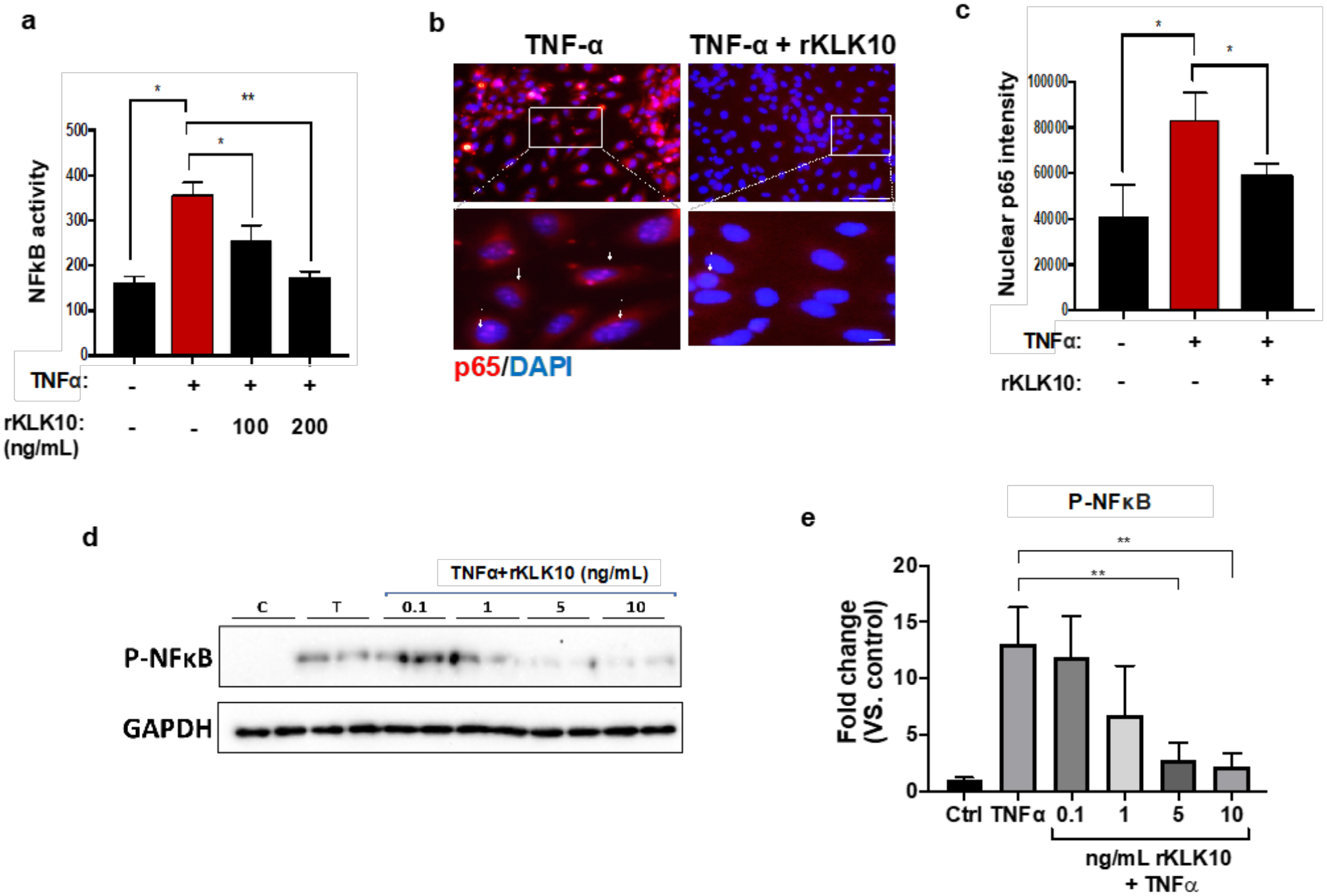
rKLK10 inhibits NFκB Activity. (a) HAECs were transfected with an NFκB luciferase reporter plasmid and treated with TNFα (5 ng/mL) for 4h followed by rKLK10. NFκB activity was measured as luciferase activity. (b) HAECs were treated with TNFα (5 ng/mL) for 4h followed by rKLK10 (100 ng/mL) and p65 immunostaining was performed with DAPI counterstain. Red=p65, Blue=DAPI. Scale bar top=50 μm, scale bar bottom=10 μm. (c) Quantification of b as measure of p65 fluorscent intensity. (d) HAECs were treated with TNFα (5 ng/mL) for 4h followed by 0.1-10 ng/mL rKLK10 and expression of p-NFκB was assessed by western blot. (e) Quantification of KLK10 signal, normalized to GAPDH and control. Data was pooled from at least three independent experiments. One-way ANOVA with Bonferroni correction for multiple comparisons where appropriate. Mean±SEM. *P≤0.05, **P≤0.01.

**Figure S6.**
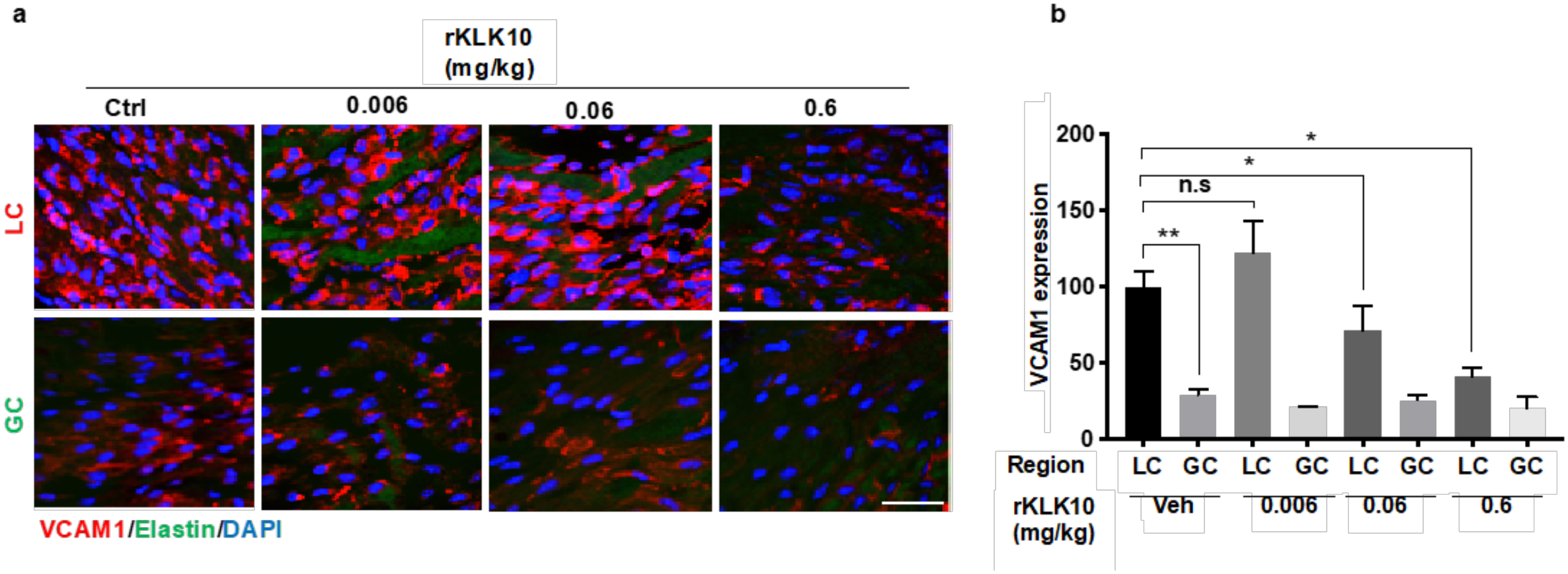
rKLK10 inhibits VCAM1 expression in the d-flow region of the mouse aortic arch in a dose-dependent manner. (a) Mice (male, C57/BL6) were administered 0.0006-0.6 mg/kg rKLK10 or vehicle by IV injection and inflammation was assessed by *en face* immunostaining of VCAM1 at the Lesser Curvature (LC) and the Greater Curvature (GC) of the aortic arch. Red=VCAM1, Blue=DAPI, Green=Elastin. Scale bar=50 um (b) Quantification of VCAM1 staining in A normalized to the LC. n=6. Two-way ANOVA with Bonferroni correction for multiple comparisons. Mean±SEM. *P≤0.05, **P ≤ 0.01.

**Figure S7.**
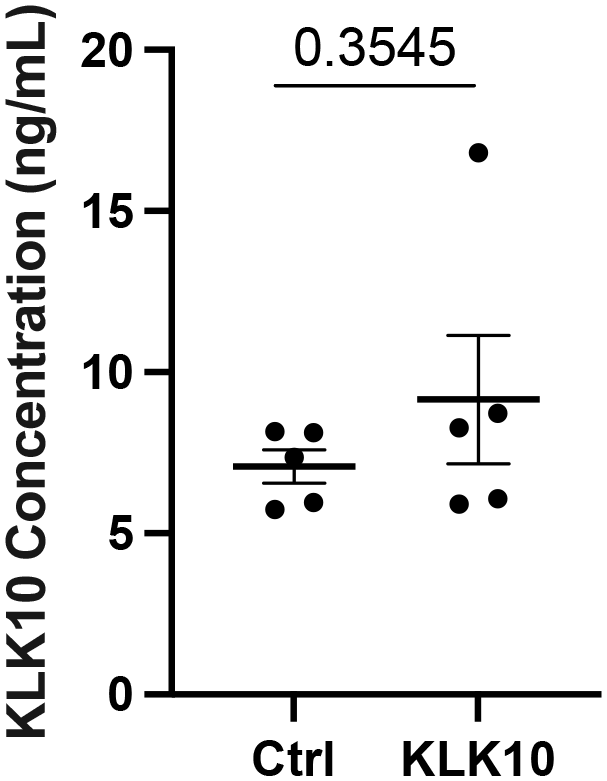
Mouse KLK10 ELISA from Ultrasound Study. KLK10 expression was measured in the plasma from mice injected with Ctrl or KLK10 plasmid and adminstered ultrasound using mouse KLK10 ELISA (Antibodies-Online ABIN628061). Paired two-tailed t-test. N=5.

**Figure S8.**
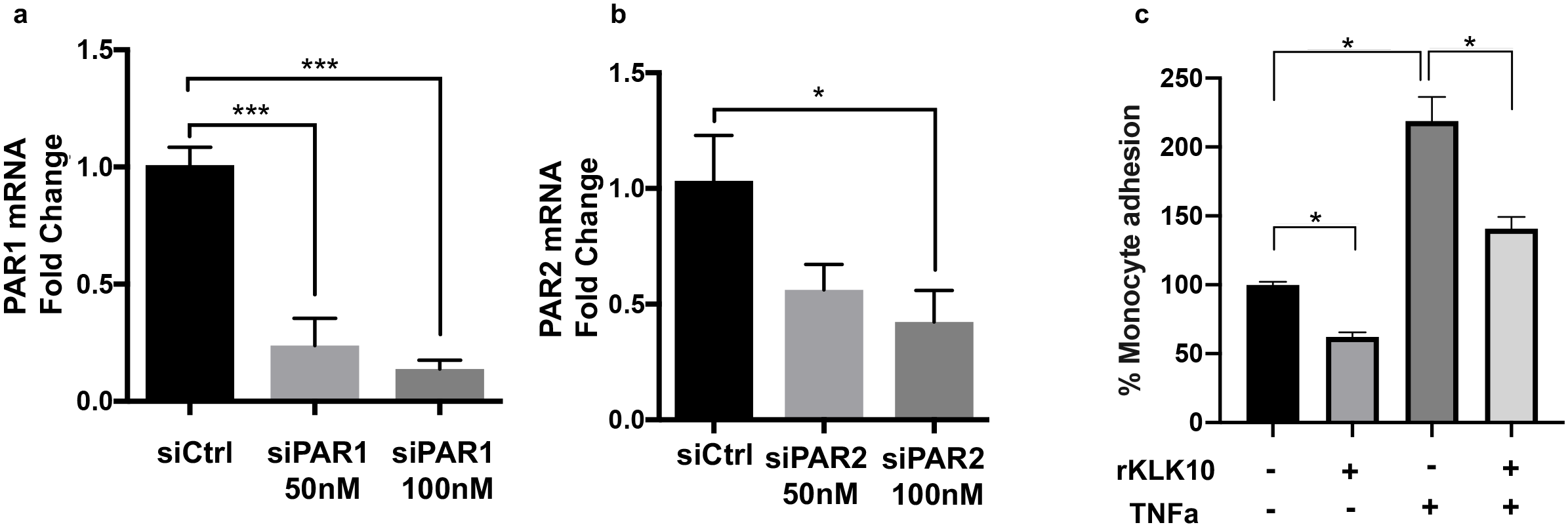
Control studies show effective knockdown of PAR1 and PAR2 with the PAR1/2 siRNAs and the anti-inflammatory effect of rKLK10. (a,b) HAECs were transfected 50 or 100nM siPAR1, siPAR2, or control siRNA (siCtrl) and expression of PAR1 and PAR2 mRNAs were measured by qPCR. (c) HAECs were treated with rKLK10 (100 ng/mL), TNFα (5 ng/mL), or rKLK10 and TNFα, and the THP1 monocyte adhesion to EC was performed in parallel with PAR1/2 SEAP assay shown in Figure 6c,d. n=4. One-way ANOVA with Bonferroni correction for multiple comparisons where appropriate. Mean±SEM. *P≤0.05, ***P ≤ 0.001.

**Figure S9.**
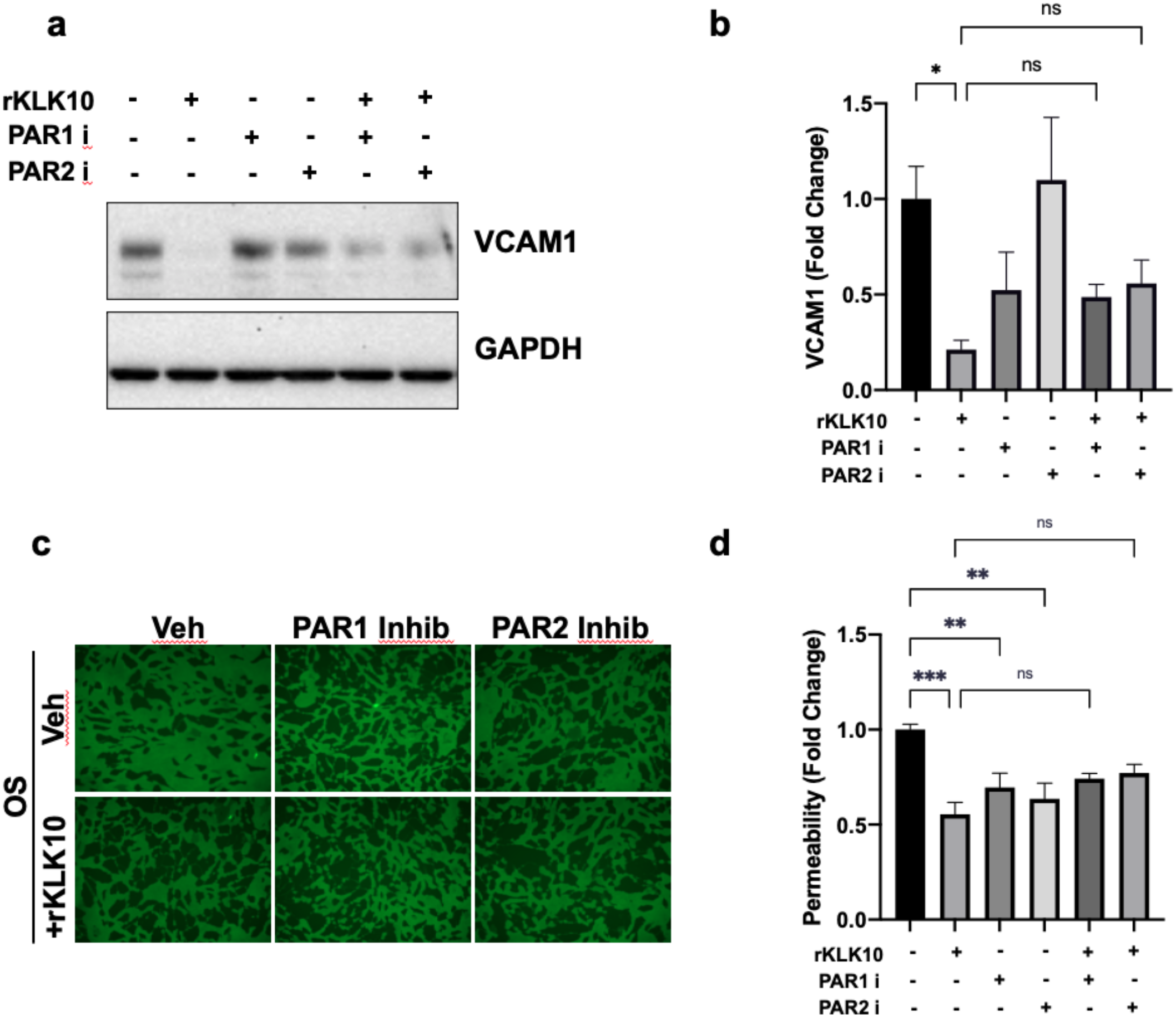
PAR1/2 inhibition in HAECs exposed to OS and treated with rKLK10. (a) HAECs were exposed to OS for 24hr with rKLK10 (10 ng/mL) and PAR1 (SCH79797, 0.1 μM) or PAR2 inhibitors ( FSLLRY-NH2, 10 μM). Cell lysate was collected for western blot analysis of VCAM1 expression. (b) Quantification of VCAM1 signal in A, normalized to GAPDH and the vehicle control. n=4. (c) HAECs were exposed to OS for 24hr with rKLK10 and PAR1/2 inhibitiors and the FITC-avidin permeability assay was performed. (d) Quantification of permeability in C, normalized to vehicle control. One-way ANOVA with Bonferroni correction for multiple comparisons where appropriate. Mean±SEM. *P≤0.05.

**Figure S10.**
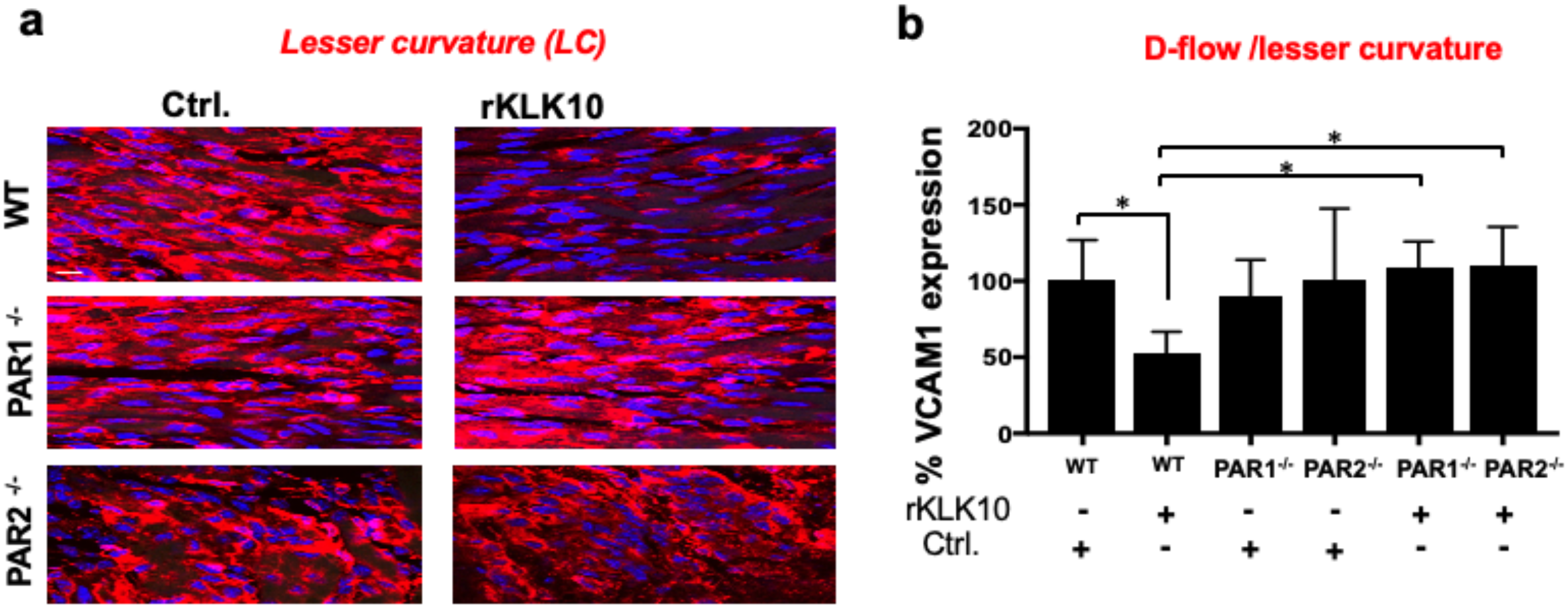
Knockdown of PAR1 or PAR2 *in vivo* attenuates the anti-inflammatory effect of KLK10 at the lesser curvature. (a) C57/BL6 wild-type, PAR1^-/-^, and PAR2^-/-^ mice were injected with rKLK10 (0.6 mg/kg once every 3 days for 1 week) and the aortic arch LC and GC were en face immunostained with the VCAM1 antibody (red) and DAPI (blue). (b) Quantification of VCAM1 fluorescence in A, normalized to the % of VCAM1 expression WT ctrl mice. n=4. One-way ANOVA with Bonferroni correction for multiple comparisons where appropriate. Mean±SEM. *P≤0.05.

**Figure S11.**
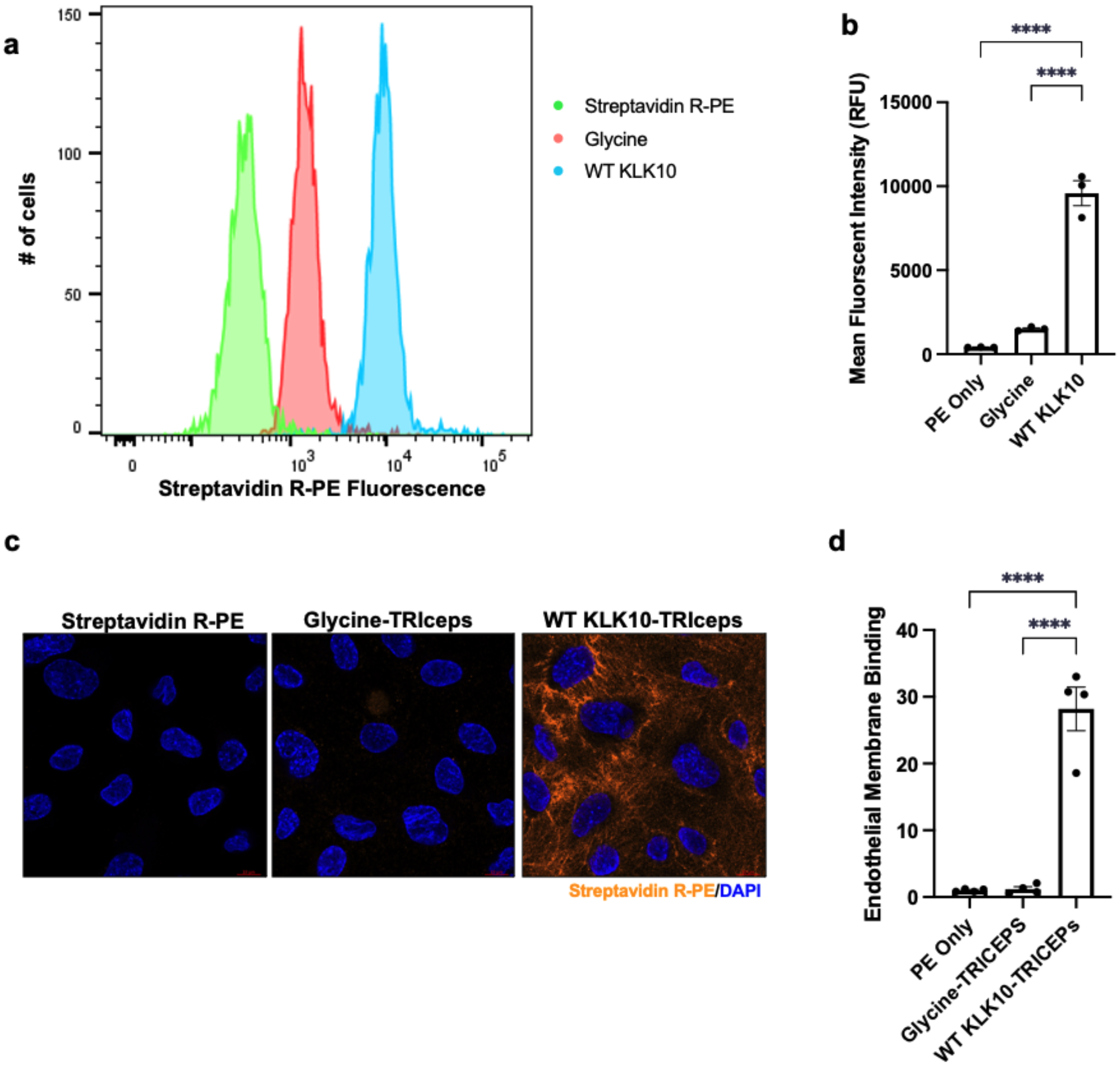
KLK10-coupled TRICEPS binds the endothelial membrane. WT rKLK10-coupled TRICEPS was added to HAECs and binding to the membrane was assessed by flow cytometry (a,b) and immunostaining (c,d) using Streptavidin-PE which binds to the biotin on TRICEPS. Glycine-coupled TRICEPS and Streptavidin-PE alone were used as a negative controls. n=3-4. One-way ANOVA with Bonferroni correction for multiple comparisons where appropriate. Mean±SEM. *P≤0.05.

**Figure S12.**
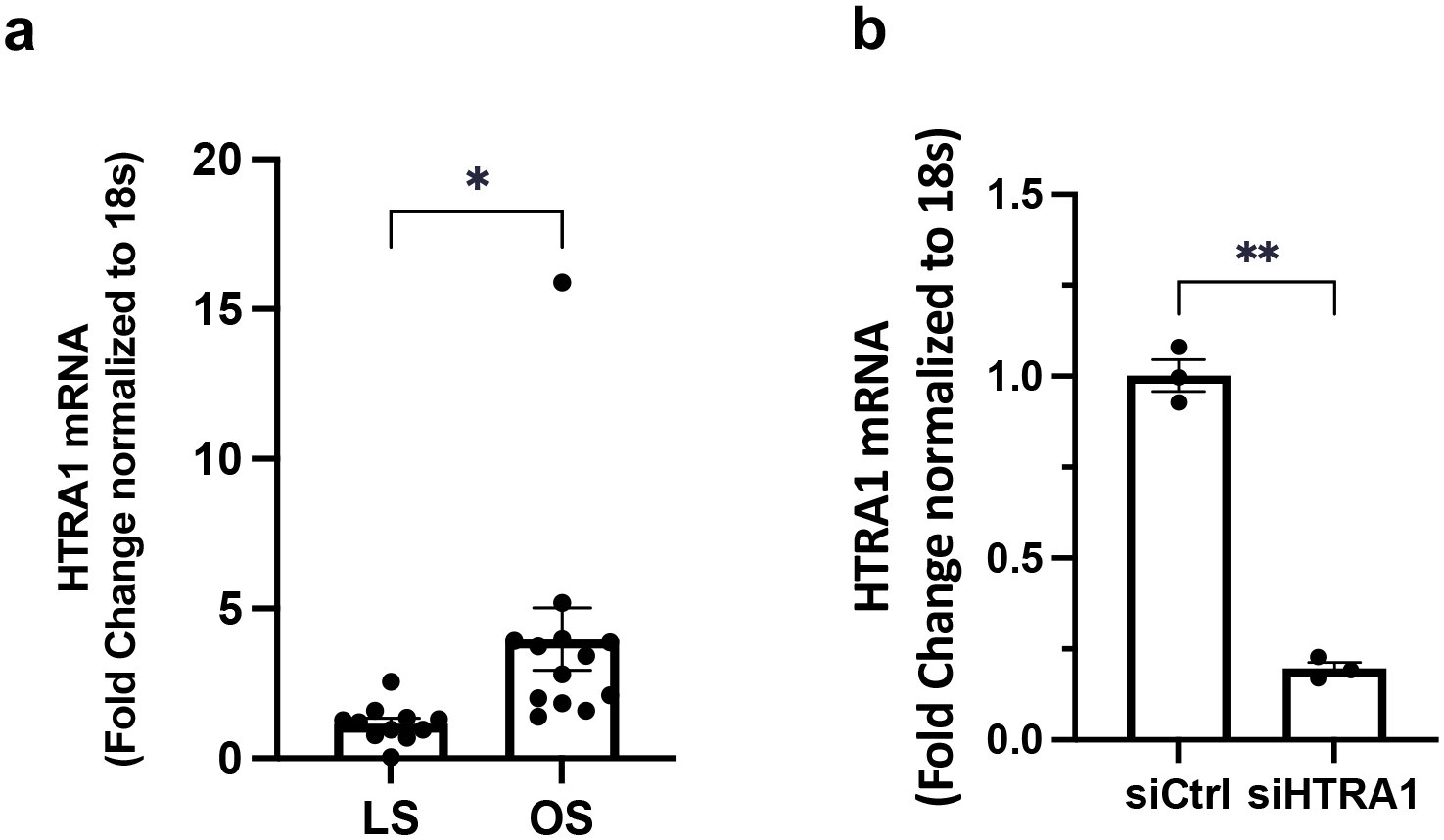
HTRA1 Shear Sensitivity and knockdown by siRNA. (a) HAECs were exposed to LS or OS for 24hrs and HTRA1 shear sensitivity was assessed by qPCR. Data is normalized to 18s and LS control. (b) HAECs were transfected with siRNA corresponding to HTRA1 for 48hrs and knockdown of HTRA1 was assessed by qPCR. Data is normalized to 18s and siCtrl. n=3-13. Paired two-tailed t-test. Mean±SEM. *P≤0.05.

**Figure S13.**
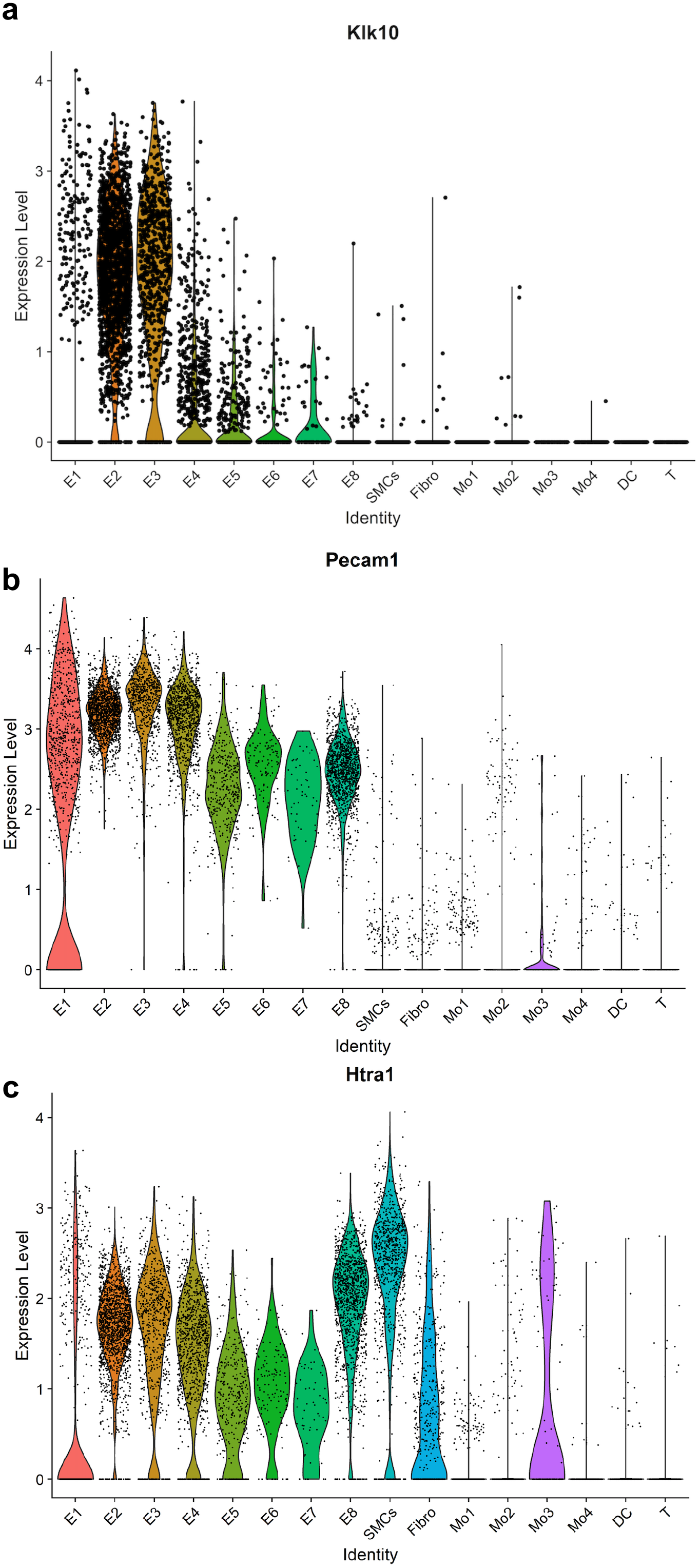
scRNAseq analysis of KLK10, CD31, and HTRA1 from the PCL mouse model. Violin plots representing single cell expression of (a) KLK10, (b) PECAM1, (c) HTRA1 genes in endothelial cells (E1-E8), Smooth Muscle Cells (SMCs), Firboblasts (Fibro), Monocytes/Macrophages (Mo1-3), Dendritic (DC), and T cells (T). E1–E4 consisted of ECs exposed to *s-flow* conditions. The majority of E5 and E7 clusters consisted of ECs exposed to acute *d-flow.* E6 and E8 exclusively consisted of ECs exposed to chronic *d-flow*.

**Figure S14.**
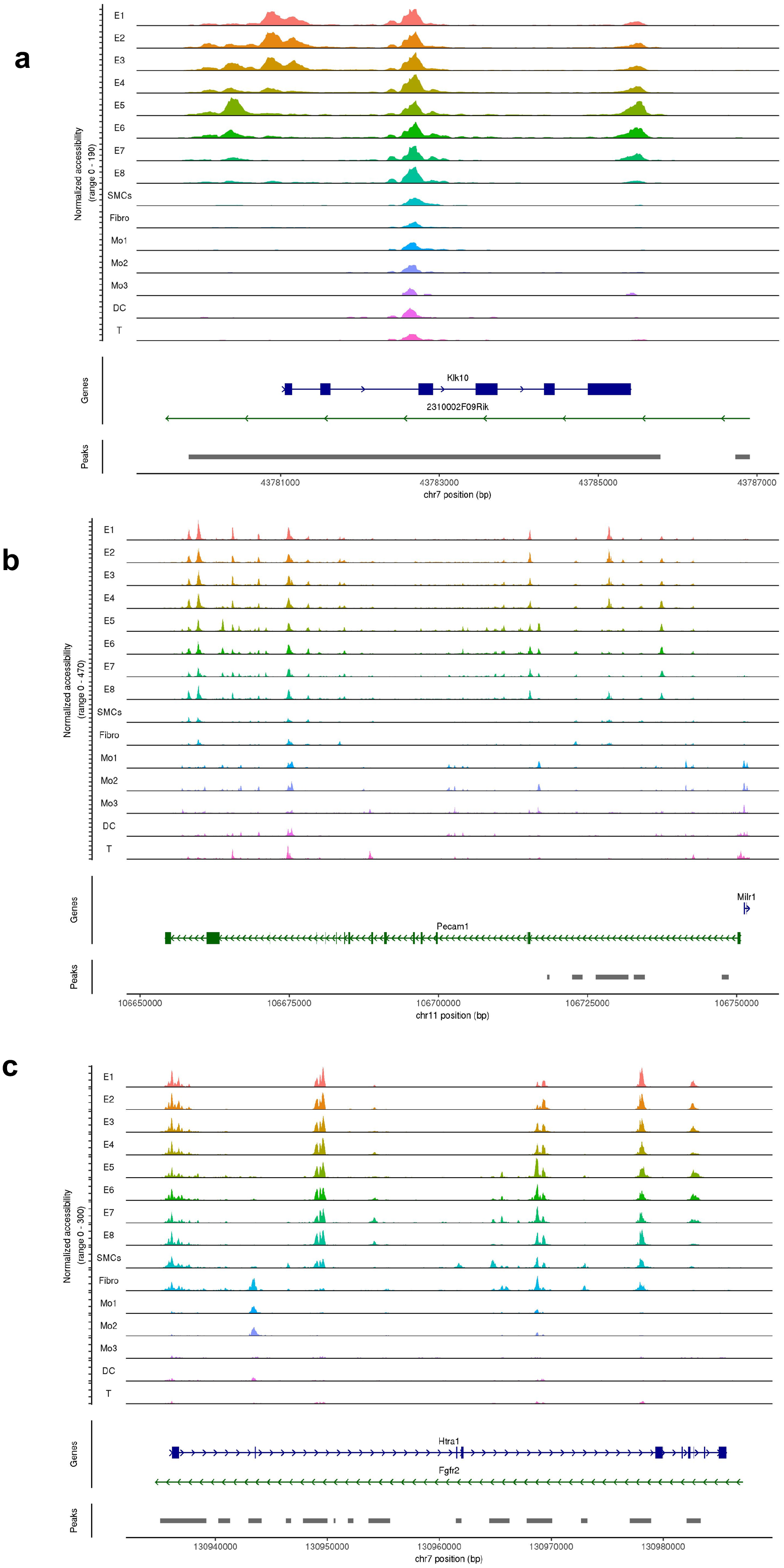
scATACseq analysis of KLK10, CD31, and HTRA1 from the PCL mouse model. Chromatin accessibility plots displaying single cell chromatin state of (a) KLK10, (b) PECAM1, (c) HTRA1 genes in endothelial cells (E1-E8), Smooth Muscle Cells (SMCs), Firboblasts (Fibro), Monocytes/Macrophages (Mo1-3), Dendritic (DC), and T cells (T). E1–E4 consisted of ECs exposed to *s-flow* conditions. The majority of E5 and E7 clusters consisted of ECs exposed to acute *d-flow.* E6 and E8 exclusively consisted of ECs exposed to chronic *d-flow*.

**Figure S15.**
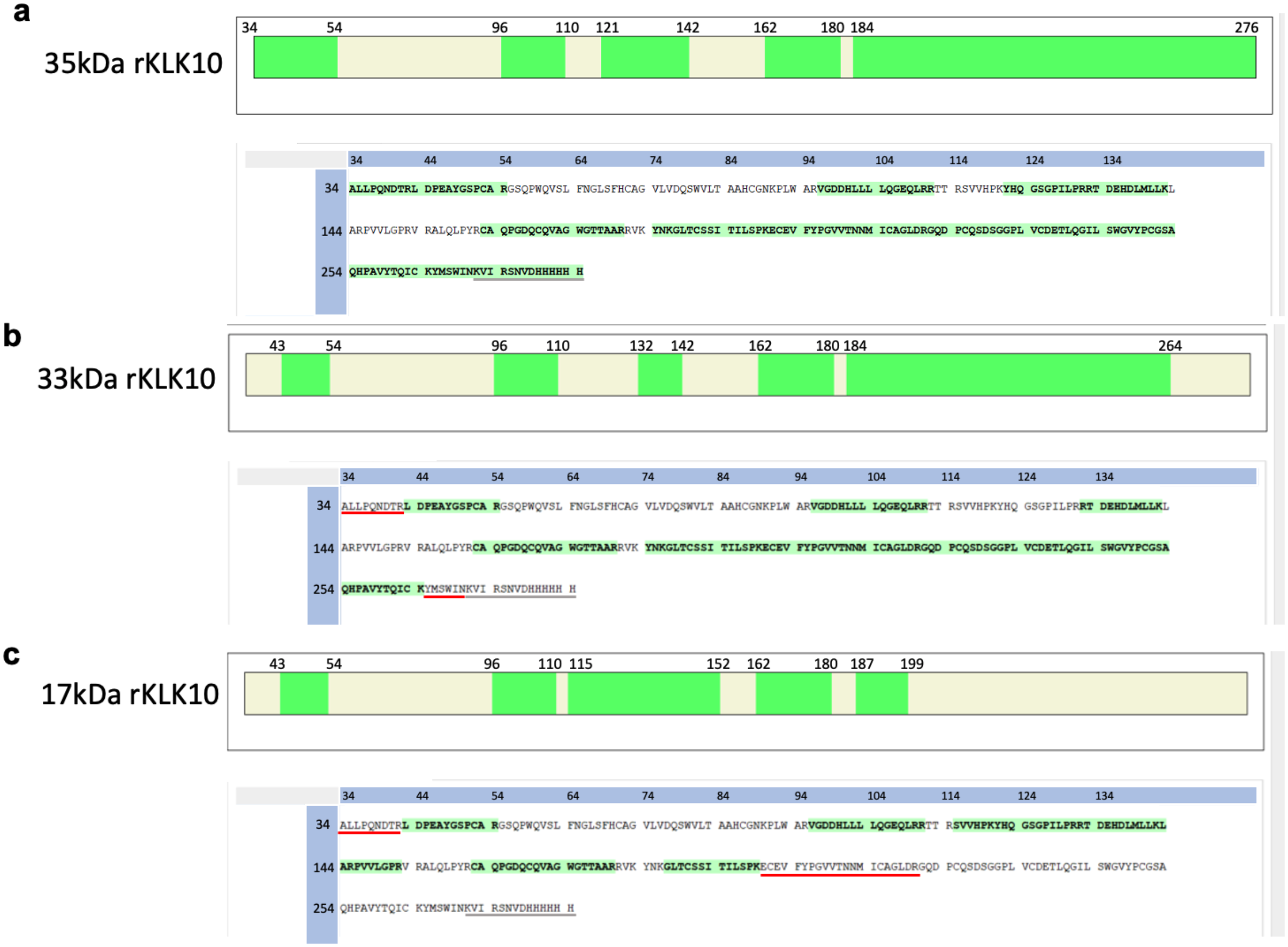
Tryptic peptide map of rKLK10 cleaved by HTRA1. rKLK10 was incubated with WT or catalytically inactive HTRA1 (500 ng each) for 18hrs and resolved by SDS-PAGE. Coomassie-stained bands corresponding to 35kDa, 33kDa, and 17kDa rKLK10 were excised and digested with trypsin followed by tandem liquid chromatography mass spectrometry. Peptide sequences were analyzed using Proteome Discoverer and were mapped against 35kDa rKLK10 (A34-N276) protein sequence. Tryptic sequences identified by the mass spectrometry are highlighted in green. Red lines indicate potential cleavage regions by HTRA1. For clarity, the signal peptide (M1-A33) is not included and the C-terminal His-tag sequence is marked with a gray line.

**Figure S16.**
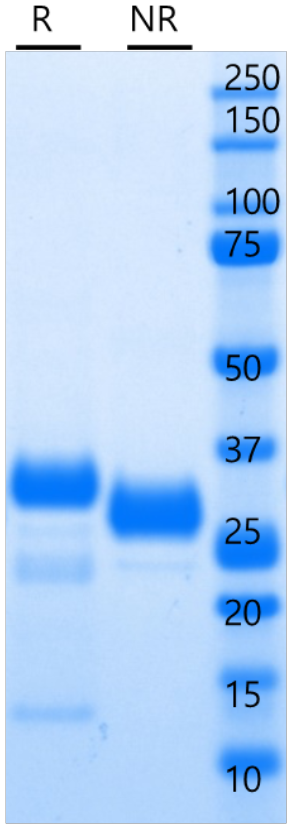
Human rKLK10 purification from CHO cells. Human rKLK10 with His-tag overexpressed in CHO cells was purified by affinity chromotagraphy. Purified rKLK10 (5 ug) was resolved by SDS-PAGE under reducing (R) or non-reducing (NR) conditions and stained with Coomasie blue.

**Table S1.**
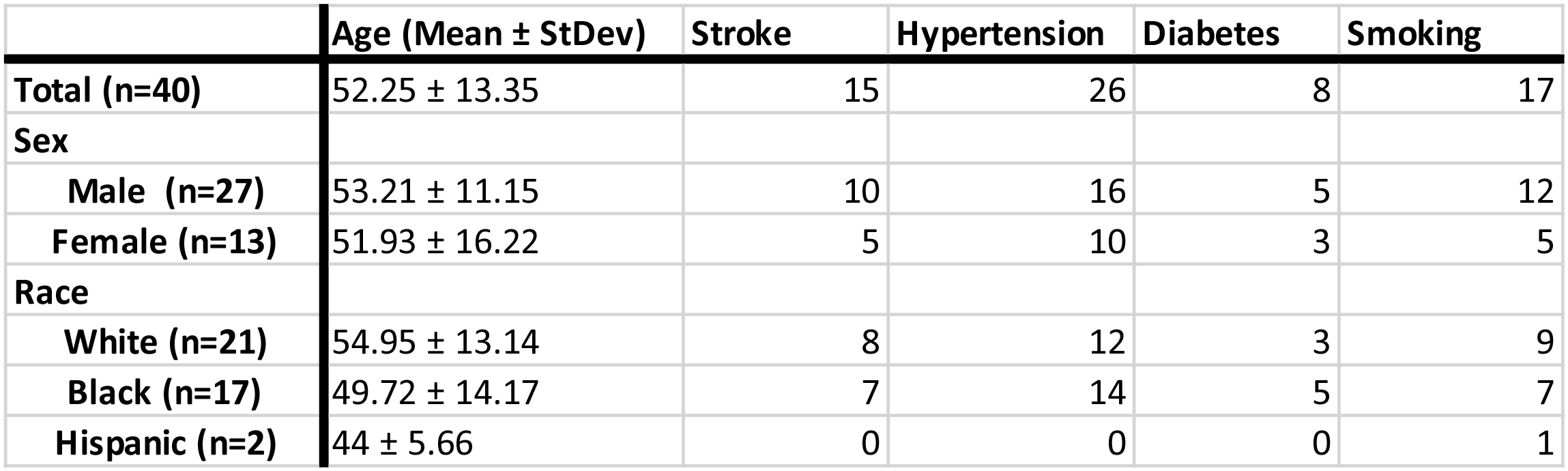
Patient Characteristics.

**Table S2.**
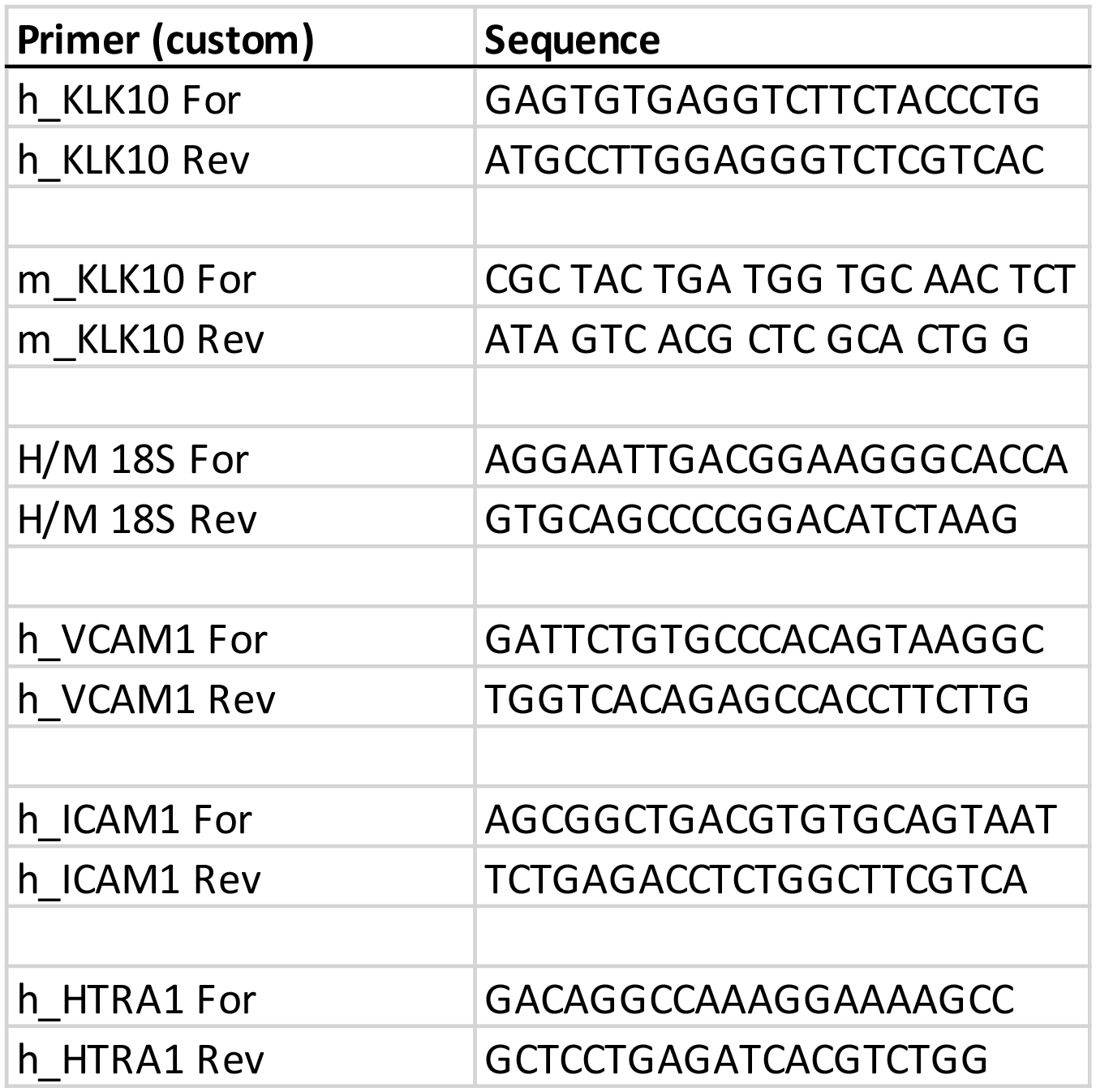
qPCR Primers.

## References

1. Chiu J-J, Chien S. Effects of disturbed flow on vascular endothelium: Pathophysiological basis and clinical perspectives. Physiol. Rev. 2011;91:327–387

2. Davies PF. Flow-mediated endothelial mechanotransduction. Physiol. Rev. 1995;75:519–560

3. Kwak BR, Back M, Bochaton-Piallat ML, Caligiuri G, Daemen MJ, Davies PF, Hoefer IE, Holvoet P, Jo H, Krams R, Lehoux S, Monaco C, Steffens S, Virmani R, Weber C, Wentzel JJ, Evans PC. Biomechanical factors in atherosclerosis: Mechanisms and clinical implications. Eur Heart J. 2014;35:3013–3020, 3020a–3020d

4. Tarbell JM, Shi ZD, Dunn J, Jo H. Fluid mechanics, arterial disease, and gene expression. Annu Rev Fluid Mech. 2014;46:591–614

5. Mack JJ, Mosqueiro TS, Archer BJ, Jones WM, Sunshine H, Faas GC, Briot A, Aragón RL, Su T, Romay MC. Notch1 is a mechanosensor in adult arteries. Nature communications. 2017;8:1–19

6. Tzima E, Irani-Tehrani M, Kiosses WB, Dejana E, Schultz DA, Engelhardt B, Cao G, DeLisser H, Schwartz MA. A mechanosensory complex that mediates the endothelial cell response to fluid shear stress. Nature. 2005;437:426–431

7. Li J, Hou B, Muraki K, Ainscough J, Beech D. Piezo1 integration of vascular architecture with physiological force. The FASEB Journal. 2015;29:639.632

8. Chachisvilis M, Zhang YL, Frangos JA. G protein-coupled receptors sense fluid shear stress in endothelial cells. Proc. Natl. Acad. Sci. U. S. A. 2006;103:15463–15468

9. Florian JA, Kosky JR, Ainslie K, Pang Z, Dull RO, Tarbell JM. Heparan sulfate proteoglycan is a mechanosensor on endothelial cells. Circ. Res. 2003;93:e136–e142

10. Wang L, Luo J-Y, Li B, Tian XY, Chen L-J, Huang Y, Liu J, Deng D, Lau CW, Wan S. Integrin-yap/taz-jnk cascade mediates atheroprotective effect of unidirectional shear flow. Nature. 2016;540:579–582

11. Kumar S, Kim CW, Simmons RD, Jo H. Role of flow-sensitive micrornas in endothelial dysfunction and atherosclerosis: Mechanosensitive athero-mirs. Arterioscler. Thromb. Vasc. Biol. 2014;34:2206–2216

12. Kumar S, Williams D, Sur S, Wang J-Y, Jo H. Role of flow-sensitive micrornas and long noncoding rnas in vascular dysfunction and atherosclerosis. Vascular Pharmacology. 2019;114:76–92

13. Dunn J, Qiu H, Kim S, Jjingo D, Hoffman R, Kim CW, Jang I, Son DJ, Kim D, Pan C, Fan Y, Jordan IK, Jo H. Flow-dependent epigenetic DNA methylation regulates endothelial gene expression and atherosclerosis. The Journal of Clinical Investigation. 2014;124:3187–3199

14. Nam D, Ni C-W, Rezvan A, Suo J, Budzyn K, Llanos A, Harrison D, Giddens D, Jo H. Partial carotid ligation is a model of acutely induced disturbed flow, leading to rapid endothelial dysfunction and atherosclerosis. American Journal of Physiology-Heart and Circulatory Physiology. 2009;297:H1535–H1543

15. Ni C-W, Qiu H, Rezvan A, Kwon K, Nam D, Son DJ, Visvader JE, Jo H. Discovery of novel mechanosensitive genes in vivo using mouse carotid artery endothelium exposed to disturbed flow. Blood. 2010;116:e66–e73

16. Diamandis EP, Yousef GM, Clements J, Ashworth LK, Yoshida S, Egelrud T, Nelson PS, Shiosaka S, Little S, Lilja H, Stenman UH, Rittenhouse HG, Wain H. New nomenclature for the human tissue kallikrein gene family. Clin Chem. 2000;46:1855–1858

17. Yousef GM, Luo LY, Diamandis EP. Identification of novel human kallikrein-like genes on chromosome 19q13.3-q13.4. Anticancer Res. 1999;19:2843–2852

18. Yousef GM, Diamandis EP. An overview of the kallikrein gene families in humans and other species: Emerging candidate tumour markers. Clin. Biochem. 2003;36:443–452

19. Madeddu P, Emanueli C, El-Dahr S. Mechanisms of disease: The tissue kallikrein–kinin system in hypertension and vascular remodeling. Nature Clinical Practice Nephrology. 2007;3:208–221

20. Clements JA, Willemsen NM, Myers SA, Dong Y. The tissue kallikrein family of serine proteases: Functional roles in human disease and potential as clinical biomarkers. Crit Rev Clin Lab Sci. 2004;41:265–312

21. Yousef GM, Diamandis EP. The new human tissue kallikrein gene family: Structure, function, and association to disease. Endocrine reviews. 2001;22:184–204

22. Pampalakis G, Sotiropoulou G. Tissue kallikrein proteolytic cascade pathways in normal physiology and cancer. Biochimica et Biophysica Acta (BBA)-Reviews on Cancer. 2007;1776:22–31

23. Margolius HS. Tissue kallikreins structure, regulation, and participation in mammalian physiology and disease. Clinical reviews in allergy & immunology. 1998;16:337–349

24. Campbell DJ. The kallikrein–kinin system in humans. Clinical and experimental pharmacology and physiology. 2001;28:1060–1065

25. Goyal J, Smith KM, Cowan JM, Wazer DE, Lee SW, Band V. The role for nes1 serine protease as a novel tumor suppressor. Cancer Res. 1998;58:4782–4786

26. Liu X-L, Wazer DE, Watanabe K, Band V. Identification of a novel serine protease-like gene, the expression of which is down-regulated during breast cancer progression. Cancer Res. 1996;56:3371–3379

27. Hu J, Lei H, Fei X, Liang S, Xu H, Qin D, Wang Y, Wu Y, Li B. Nes1/klk10 gene represses proliferation, enhances apoptosis and down-regulates glucose metabolism of pc3 prostate cancer cells. Sci. Rep. 2015;5:17426

28. Luo LY, Rajpert-De Meyts ER, Jung K, Diamandis EP. Expression of the normal epithelial cell-specific 1 (nes1; klk10) candidate tumour suppressor gene in normal and malignant testicular tissue. Br J Cancer. 2001;85:220–224

29. Zhang Y, Song H, Miao Y, Wang R, Chen L. Frequent transcriptional inactivation of kallikrein 10 gene by cpg island hypermethylation in non-small cell lung cancer. Cancer Sci. 2010;101:934–940

30. Luo L-Y, Katsaros D, Scorilas A, Fracchioli S, Bellino R, van Gramberen M, de Bruijn H, Henrik A, Stenman U-H, Massobrio M, van der Zee AGJ, Vergote I, Diamandis EP. The serum concentration of human kallikrein 10 represents a novel biomarker for ovarian cancer diagnosis and prognosis. Cancer Res. 2003;63:807–811

31. Yousef GM, White NM, Michael IP, Cho JC, Robb JD, Kurlender L, Khan S, Diamandis EP. Identification of new splice variants and differential expression of the human kallikrein 10 gene, a candidate cancer biomarker. Tumour Biol. 2005;26:227–235

32. Dorn J, Bayani J, Yousef GM, Yang F, Magdolen V, Kiechle M, Diamandis EP, Schmitt M. Clinical utility of kallikrein-related peptidases (klk) in urogenital malignancies. Thromb. Haemost. 2013;110:408–422

33. Tailor PD, Kodeboyina SK, Bai S, Patel N, Sharma S, Ratnani A, Copland JA, She J-X, Sharma A. Diagnostic and prognostic biomarker potential of kallikrein family genes in different cancer types. Oncotarget. 2018;9

34. Sidiropoulos M, Pampalakis G, Sotiropoulou G, Katsaros D, Diamandis EP. Downregulation of human kallikrein 10 (klk10/nes1) by cpg island hypermethylation in breast, ovarian and prostate cancers. Tumour Biol. 2005;26:324–336

35. Bharaj BB, Luo LY, Jung K, Stephan C, Diamandis EP. Identification of single nucleotide polymorphisms in the human kallikrein 10 (klk10) gene and their association with prostate, breast, testicular, and ovarian cancers. The Prostate. 2002;51:35–41

36. Jo H, Song H, Mowbray A. Role of nadph oxidases in disturbed flow-and bmp4-induced inflammation and atherosclerosis. Antioxidants & redox signaling. 2006;8:1609–1619

37. Chang K, Weiss D, Suo J, Vega JD, Giddens D, Taylor WR, Jo H. Bone morphogenic protein antagonists are coexpressed with bone morphogenic protein 4 in endothelial cells exposed to unstable flow in vitro in mouse aortas and in human coronary arteries. Circulation. 2007;116:1258–1266

38. Andueza A, Kumar S, Kim J, Kang D-W, Mumme HL, Perez JI, Villa-Roel N, Jo H. Endothelial reprogramming by disturbed flow revealed by single-cell rna and chromatin accessibility study. Cell reports. 2020;33:108491

39. Dubrovskyi O, Birukova AA, Birukov KG. Measurement of local permeability at subcellular level in cell models of agonist- and ventilator-induced lung injury. Laboratory investigation; a journal of technical methods and pathology. 2013;93:254–263

40. Liu R, Tang J, Xu Y, Dai Z. Bioluminescence imaging of inflammation in vivo based on bioluminescence and fluorescence resonance energy transfer using nanobubble ultrasound contrast agent. ACS Nano. 2019;13:5124–5132

41. Borden MA, Kruse DE, Caskey CF, Zhao S, Dayton PA, Ferrara KW. Influence of lipid shell physicochemical properties on ultrasound-induced microbubble destruction. IEEE Trans Ultrason Ferroelectr Freq Control. 2005;52:1992–2002

42. Shapiro G, Wong AW, Bez M, Yang F, Tam S, Even L, Sheyn D, Ben-David S, Tawackoli W, Pelled G, Ferrara KW, Gazit D. Multiparameter evaluation of in vivo gene delivery using ultrasound-guided, microbubble-enhanced sonoporation. J Control Release. 2016;223:157–164

43. Oikonomopoulou K, Hansen KK, Saifeddine M, Tea I, Blaber M, Blaber SI, Scarisbrick I, Andrade-Gordon P, Cottrell GS, Bunnett NW, Diamandis EP, Hollenberg MD. Proteinase-activated receptors, targets for kallikrein signaling. J Biol Chem. 2006;281:32095–32112

44. Oikonomopoulou K, Hansen KK, Saifeddine M, Vergnolle N, Tea I, Blaber M, Blaber SI, Scarisbrick I, Diamandis EP, Hollenberg MD. Kallikrein-mediated cell signalling: Targeting proteinase-activated receptors (pars). Biol Chem. 2006;387:817–824

45. Coughlin SR. Protease-activated receptors in hemostasis, thrombosis and vascular biology. J Thromb Haemost. 2005;3:1800–1814

46. Coughlin SR. Thrombin signalling and protease-activated receptors. Nature. 2000;407:258–264

47. Rana S, Yang L, Hassanian SM, Rezaie AR. Determinants of the specificity of protease-activated receptors 1 and 2 signaling by factor xa and thrombin. J Cell Biochem. 2012;113:977–984

48. Mosnier LO, Sinha RK, Burnier L, Bouwens EA, Griffin JH. Biased agonism of protease-activated receptor 1 by activated protein c caused by noncanonical cleavage at arg46. Blood. 2012;120:5237–5246

49. Bae JS, Yang L, Rezaie AR. Lipid raft localization regulates the cleavage specificity of protease activated receptor 1 in endothelial cells. J. Thromb. Haemost. 2008;6:954–961

50. Frei AP, Jeon O-Y, Kilcher S, Moest H, Henning LM, Jost C, Plückthun A, Mercer J, Aebersold R, Carreira EM. Direct identification of ligand-receptor interactions on living cells and tissues. Nat. Biotechnol. 2012;30:997

51. Frei AP, Moest H, Novy K, Wollscheid B. Ligand-based receptor identification on living cells and tissues using triceps. Nat. Protoc. 2013;8:1321–1336

52. Chen CY, Melo E, Jakob P, Friedlein A, Elsässer B, Goettig P, Kueppers V, Delobel F, Stucki C, Dunkley T, Fauser S, Schilling O, Iacone R. N-terminomics identifies htra1 cleavage of thrombospondin-1 with generation of a proangiogenic fragment in the polarized retinal pigment epithelial cell model of age-related macular degeneration. Matrix Biol. 2018;70:84–101

53. Jiang J, Huang L, Yu W, Wu X, Zhou P, Li X. Overexpression of htra1 leads to down-regulation of fibronectin and functional changes in rf/6a cells and huvecs. PLoS One. 2012;7:e46115–e46115

54. Launay S, Maubert E, Lebeurrier N, Tennstaedt A, Campioni M, Docagne F, Gabriel C, Dauphinot L, Potier MC, Ehrmann M, Baldi A, Vivien D. Htra1-dependent proteolysis of tgf-beta controls both neuronal maturation and developmental survival. Cell Death Differ. 2008;15:1408–1416

55. Oka C, Tsujimoto R, Kajikawa M, Koshiba-Takeuchi K, Ina J, Yano M, Tsuchiya A, Ueta Y, Soma A, Kanda H, Matsumoto M, Kawaichi M. Htra1 serine protease inhibits signaling mediated by tgfbeta family proteins. Development. 2004;131:1041–1053

56. Son DJ, Kumar S, Takabe W, Woo Kim C, Ni C-W, Alberts-Grill N, Jang I-H, Kim S, Kim W, Won Kang S, Baker AH, Woong Seo J, Ferrara KW, Jo H. The atypical mechanosensitive microrna-712 derived from pre-ribosomal rna induces endothelial inflammation and atherosclerosis. Nature Communications. 2013;4:3000

57. Kim CW, Song H, Kumar S, Nam D, Kwon HS, Chang KH, Son DJ, Kang D-W, Brodie SA, Weiss D, Vega JD, Alberts-Grill N, Griendling K, Taylor WR, Jo H. Anti-inflammatory and antiatherogenic role of bmp receptor ii in endothelial cells. Arterioscler. Thromb. Vasc. Biol. 2013;33:1350–1359

58. Ni C-W, Qiu H, Jo H. Microrna-663 upregulated by oscillatory shear stress plays a role in inflammatory response of endothelial cells. American Journal of Physiology-Heart and Circulatory Physiology. 2011;300:H1762–H1769

59. Tressel SL, Huang RP, Tomsen N, Jo H. Laminar shear inhibits tubule formation and migration of endothelial cells by an angiopoietin-2 dependent mechanism. Arterioscler Thromb Vasc Biol. 2007;27:2150–2156

60. Alberts-Grill N, Rezvan A, Son DJ, Qiu H, Kim CW, Kemp ML, Weyand CM, Jo H. Dynamic immune cell accumulation during flow-induced atherogenesis in mouse carotid artery: An expanded flow cytometry method. Arteriosclerosis, thrombosis, and vascular biology. 2012;32:623–632

61. Wang Y, Sun H-Y, Kumar S, Puerta MdM, Jo H, Rezvan A. Zbtb46 is a shear-sensitive transcription factor inhibiting endothelial cell proliferation via gene expression regulation of cell cycle proteins. Laboratory Investigation. 2019;99:305–318

62. Schägger H. Tricine–sds-page. Nat. Protoc. 2006;1:16–22

63. Wingo TS, Duong DM, Zhou M, Dammer EB, Wu H, Cutler DJ, Lah JJ, Levey AI, Seyfried NT. Integrating next-generation genomic sequencing and mass spectrometry to estimate allele-specific protein abundance in human brain. J. Proteome Res. 2017;16:3336–3347

